# Comparing single cell datasets using DensityMorph

**DOI:** 10.1101/2021.10.28.466371

**Authors:** Kristen Feher

## Abstract

The proliferation of single cell datasets has brought a wealth of information, but also great challenges in data analysis. Obtaining a cohesive overview of multiple single cell samples is difficult and requires consideration of cell population structure - which may or may not be well defined - along with subtle shifts in expression within cell populations across samples, and changes in population frequency across samples. Ideally, all this would be integrated with the experimental design, e.g. time point, genotype, treatment etc. Data visualisation is the most effective way of communicating analysis but often this takes the form of a plethora of TSNE plots, colour coded according to marker and sample. In this manuscript, I introduce a novel exploratory data analysis and visualisation method that is centred around a novel quasi-distance (DensityMorph) between single cell samples. DensityMorph makes it possible to plot single cell samples in a manner analogous to performing principal component analysis on microarray samples. Biological interpretation is ensured by the introduction of Explanatory Components, which show how marker expression and coexpression drive the differences between samples. This method is a breakthrough in terms of displaying the most pertinent biological changes across single cell samples in a compact plot. Finally, it can be used either as a stand-alone method or to structure other types of analysis such as manual flow cytometry gating or cell population clustering.

## 2 Introduction

Bulk gene expression measurements such as microarrays represented a new era in high-throughput biology. There was a conceptually easy connection between feature (gene) and sample, in the sense that a group of samples could be represented as a **point cloud** in the multivariate feature space, and distance between samples had an intuitive meaning. Given the samples could be organised in single matrix, the standard rules of multivariate statistics applied - ok, not quite, as suddenly the curse of dimensionality had to be considered. However, it was quite easy to imagine the conceptual possibilities even if the mathematical details had to be worked out. It was possible create ordinations (e.g. PCA) of either genes or samples, and inspect how the samples or genes (resp.) contributed to the ordination with biplots. It was possible to cluster genes or samples, or go one step further and perform biclustering. If covariates were available, it was possible to perform regression or classification over the samples (linear or non-linear), or perform gene set enrichment analysis over the genes (an overview of these techniques can be found in [1]).

These days, single cell RNAseq (scRNAseq) is the predominant mode of gene expression measurement, where sampling occurs on two scales: patient, mouse etc (macro) and single cell (micro). **From now, a macro-sample will simply be referred to as <u>sample</u>**, and a micro-sample will be referred to as a single cell. The straighforward connection between feature and sample has now been lost, as each sample is itself a point cloud of single cells in a multivariate space defined by the measured biomarkers, and distance between samples is no longer a simple calculation. On the other hand, flow cytometry has been around for a long time, and there, enumeration of cell populations is the dominant mode of data analysis, where a cell population is a sub-group of single cells with similar expression profiles. This way of thinking also appears to have become dominant in scRNAseq data analysis. Often, multiple scRNAseq samples are pooled into a single dataset, the pooled dataset is clustered over the single cells, and the samples can be characterised by cluster membership counts.

In general, any given sample will have a complex multivariate structure. While clustering is a valid and intuitive technique for summarising a multivariate dataset, it cannot capture the full multivariate complexity, e.g. shape of cluster, density gradient within a cluster, correlation structure of genes within and between clusters, quality of separation from other clusters, arrangement of clusters in relation to each other. When the cell-population paradigm is applied to multiple samples, more problems occur. For example, the same population type may undergo slight shifts in expression and could lead to a blurring or masking effect when clustering a pooled dataset. On the other hand, if samples are clustered individually, then a set of rules is required to match up cell populations of the same type across samples. A further issue is that a sample is not necessarily ‘clusterable’, in the sense of having discrete clusters with inter-cluster spacing being much larger than intra-cluster spacing. **In summary, a cell-population centric analysis has the potential to hide nuanced shifts in expression**.

To overcome some of these difficulties, I have developed a sample-oriented data analysis method that is reminiscent of microarray analysis. It is ideal for the exploratory data analysis of large groups of samples with a complex experimental designs. It is based on a novel quasi-distance between samples and it therefore takes the samples’ multivariate structure into account, without relying on clustering. As such, it can detect when differences amongst a group of samples are driven by subtle expression and/or frequency shifts. This method calculates a sample-sample distance matrix which can be used to construct a point cloud of samples in a **latent space** by using Principal Coordinates Analysis (PCO) [3]. To help with interpretation, explanatory components are derived by extracting univariate quantities from the samples, which are overlaid with the sample point cloud by using partial least squares (PLS) [4]. This method does not replace cell population clustering but rather it can help to systematically interpret the clustering results of many samples. **Section 3** explains the rationale for and the concept behind DensityMorph, hopefully in a user-friendly way. **Section 4** demonstrates how to use DensityMorph on a large number of flow cytometry samples. The figures for **Sections 3 and 4** are found in **Section 6**. A discussion and outlook is found in **Section 5**. Technical details can be found in **Section 8**.

## 3 Comparisons between multivariate distributions: introducing DensityMorph

### 3.1 Concept

The central concept of this manuscript is graphically summarised in **Figure 1**. In order to gain a cohesive overview of a large number of samples, it would be desirable to arrange them as a point cloud in a latent space. They would be arranged such that neighbouring samples are similar to each other and distant samples are different from each other. By matching up the sample point cloud with the experimental design and any covariates, it would enable you to judge, for example:

- whether or not there are clusters of samples, or outlying samples,
- whether samples are aligned in specific directions or they lie in a diffuse cloud,
- whether sample groups are overlapping with each other or distinct (where sample groups could be defined by experimental design such as sex, treatment etc),
- whether there are directions that are aligned with covariates such as age,
- whether sample groups are similar or different to each other in unexpected ways.

**Figure 1:**
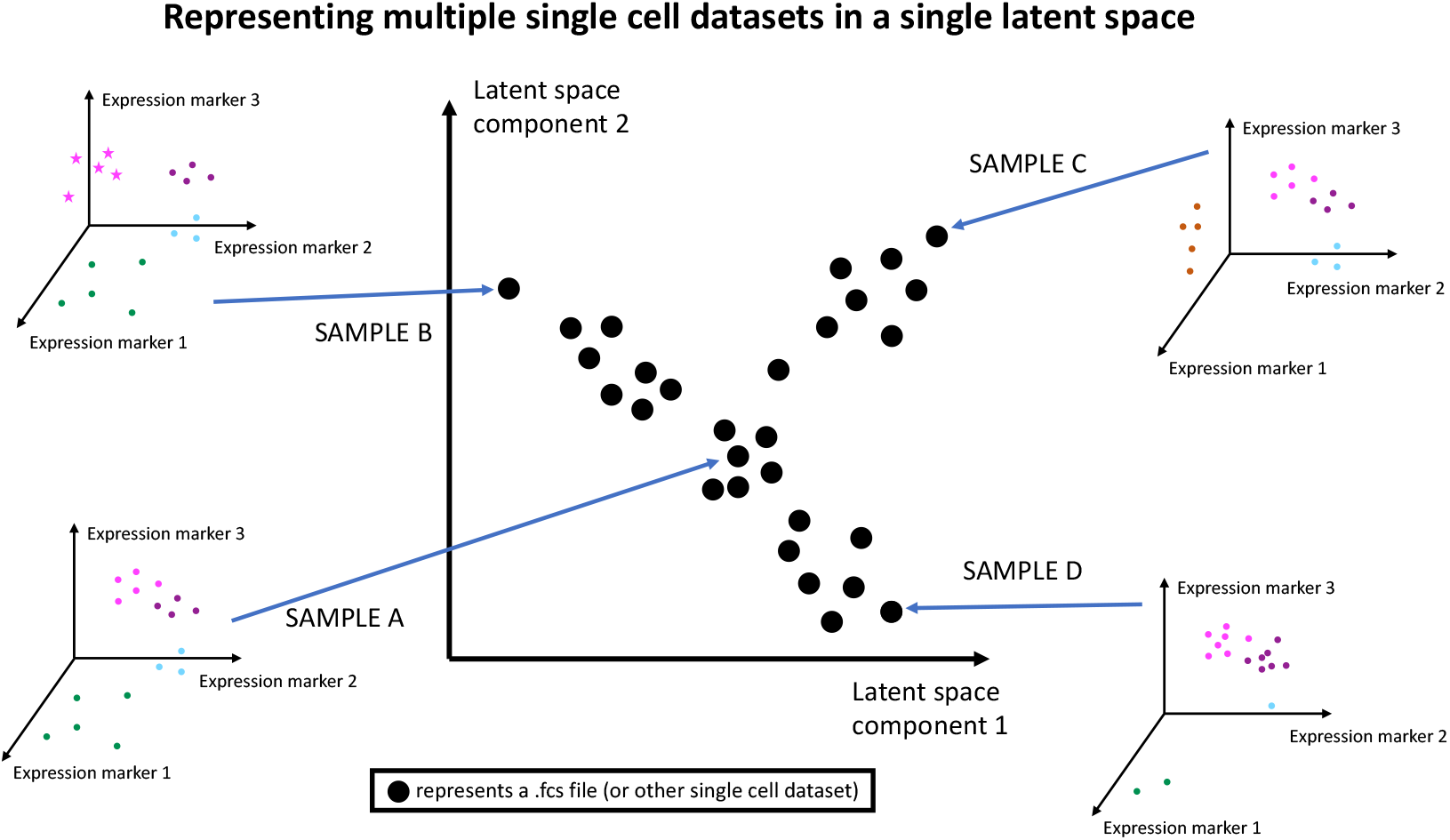
A diagram introducing the concept of embedding multiple single cell samples in a latent space, where each point in the latent space represents an entire single cell dataset. The arrangement should reflect similarity between samples, i.e. samples that are almost the same in expression and frequency patterns should be close neighbours in the latent space. In this example, sample A is at the centre of the latent space point cloud, and sample A is defined by four cell populations (green, cyan, magenta, purple) in an expression space defined by three markers. The point cloud displays 3 major directions emanating away from sample A, and the extremes are defined by samples B, C and D. They each have an overlap to sample A but cell populations shifted in expression and/or frequency, or a new population has appeared (Sample C). Samples B, C and D also differ from each other, hence they are arranged in a triangle.

In **Figure 1**, the large black point cloud represents a number of samples and Sample A, at the centre of the cloud, is like an ‘average’ sample. The cloud varies in three distinct directions, i.e. there are three continuums from Sample A to the three extremes B, C and D. They differ from A by changing the frequency of the cell populations, by shifting the cell populations, or by the presence/absence of cell populations. The sample point cloud is not a round diffuse cloud, i.e. transitional samples between B, C and D are ‘forbidden’. In a real life scenario, this could point three specific biological processes taking place, e.g. three experimental treatments on a disease, or the immune response to three different antigens. To obtain such a sample point cloud, it is necessary to measure the distance between the samples, which in turn are also point clouds.

### 3.2 Obstacles to using Wasserstein Distance

An established method of calculating distances between point clouds is the Wasserstein Distance (WD) [2]. In the simplest terms, if you imagine point clouds as piles of dirt, WD measures the energy it takes to transform one point cloud into another (it is also known as transport distance and earth mover’s distance). It is quite simple to calculate the WD in one dimension, i.e. comparing two sets of points on a line, as it is based on ranking real numbers. For *≥* 1 dimensions it becomes much harder as there is no straightforward concept of multivariate ranking. As a result, the computational time of *p*-dimensional WD (*p >* 1) is up to four orders of magnitude longer than 1-dimensional WD (**Figure 13**), rendering it impractical for routine use.

### 3.3 Summary of DensityMorph algorithm

To combat this extreme computational time, I have developed a novel approximation named ‘DensityMorph’ that compares point clouds via nearest neighbour (NN) and cross NN distances. This yields two sets of univariate points that can be used with 1-dimensional WD. Paired Density Logratios (PDLR) are the basis of DensityMorph and they are constructed using NN distances (**Figure 2**). For each point in the reference cloud (cyan), the NN distance within its own cloud is found as well as that in the comparison cloud (magenta). The logratio of the two distances signifies whether the selected point is closer to a point either in its own cloud or the other cloud. If the two point clouds have been drawn from the same underlying multivariate distribution, then the distribution of PDLR values should be symmetrically centred around zero. If this is not the case, then the comparison cloud must be different to the reference cloud.

**Figure 2:**
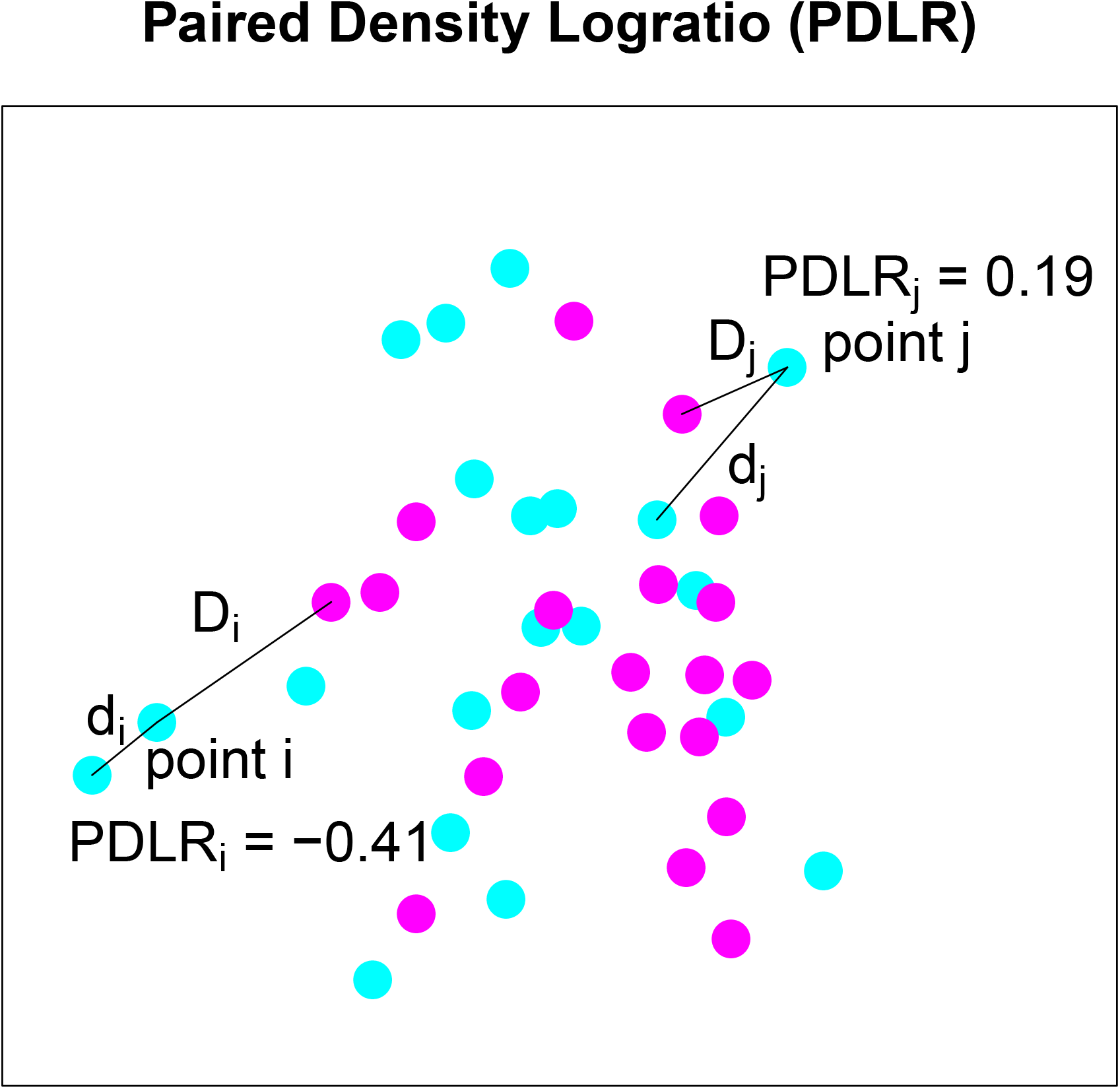
A diagram to explain the concept of Paired Density Logratio (PDLR). Two point clouds *X*_1_ (cyan) and *X*_2_ (magenta) are drawn from a bivariate normal distribution with fixed mean and variance, correlation of zero, and *N* = 20. For any given point *i* in *X*_1_, *d*_*i*_ is the distance to the nearest neighbour in *X*_1_, *D*_*i*_ is the distance to the nearest neighbour in *X*_2_. The PDLR for *i* is defined as log_10_(*d*_*i*_*/D*_*i*_). For the points selected in the diagram, PDLR_*i*_ = *−*0.41, meaning *i* is closer to another point from *X*_1_ than any point from *X*_2_. PDLR_*j*_ = 0.24, meaning *j* is closer to another point from *X*_2_ than any point from *X*_1_. The set of all PDLR values for 1…*N* is a univariate density that captures the similarity between *X*_1_ and *X*_2_. Although the concept has been demonstrated on two sets of points drawn from the same multivariate distribution, the calculation is the same for any arbitrary pair of point sets of the same dimension and *N*.

Regardless of a multivariate distribution’s shape, it seems that when comparing two point clouds drawn from the same multivariate distribution, the distribution of PDLR values is always the same (self-PDLR distribution). This fact is used to construct the DensityMorph algorithm (**Figure 3**), which approximates the distance between single cell samples such as flow cytometry samples. For every pair of samples, the cross-PDLR distribution is calculated, and the distance between the self- and cross-PDLR distributions is calculated using the 1-dimensional WD. This appears to be a good approximation for the *p*-dimensional WD and this is benchmarked in **Section 8**.

**Figure 3:**
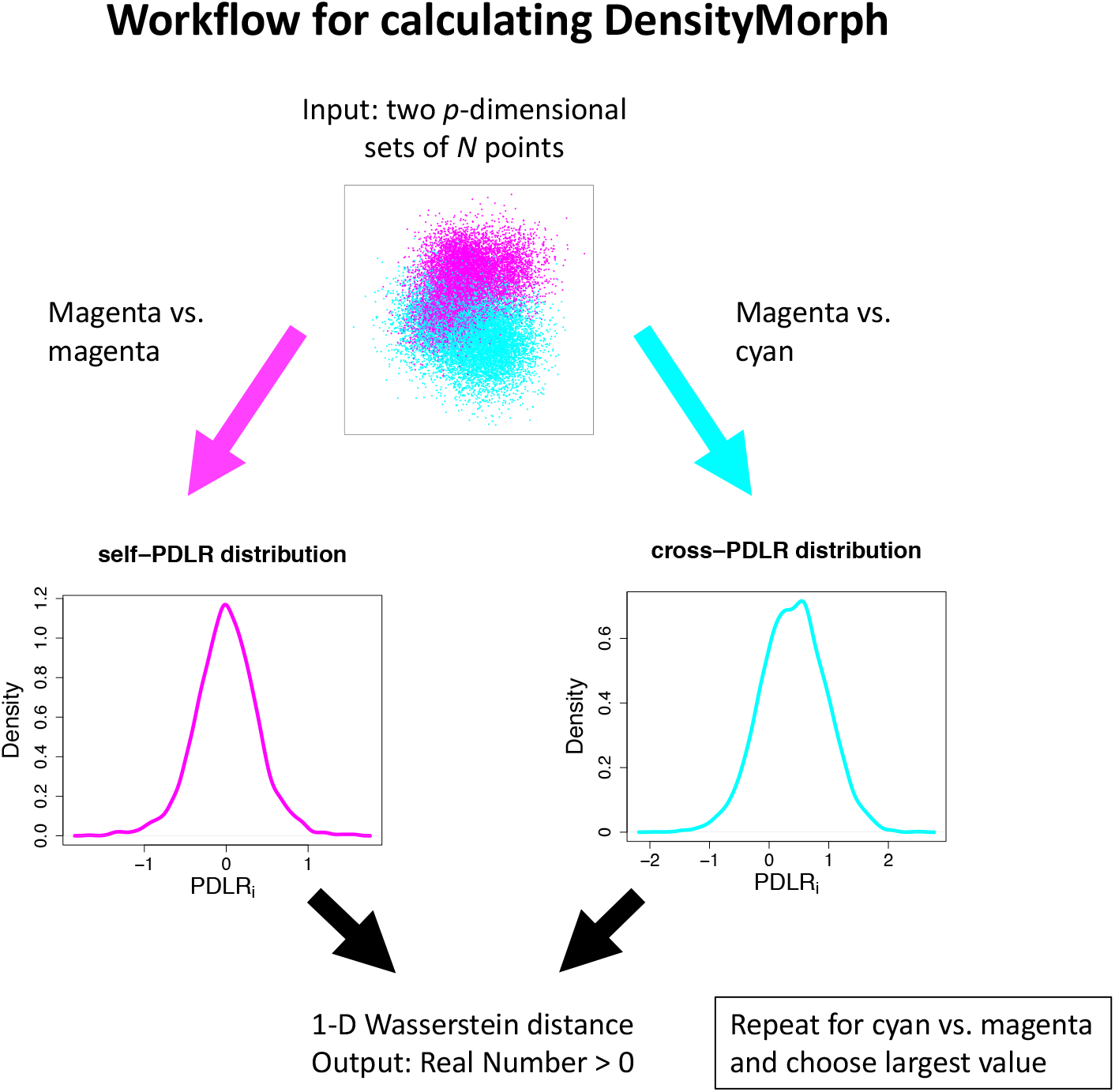
A diagram to explain the concept of DensityMorph. Two point clouds *X*_1_ (cyan) and *X*_2_ (magenta) have arbitrary *p*-dimensional distributions and *N*_1_ = *N*_2_. Following Figure 2, the self and cross PDLR distributions are calculated for *X*_1_ *→ X*_2_ and the 1-d wasserstein distance is calculated using these two distributions. The procedure is repeated for *X*_2_ *→ X*_1_ and the maximum value is chosen.

The DensityMorph algorithm can be used for characterising a set of *N* single cell samples by calculating an *N × N* distance matrix, and taking the square root of the matrix entries (**Section 8**). Principal coordinate analysis[3] is applied to this matrix yielding a set of *N* coordinates which is the sample point cloud. The latent space components (LSC) have no intrinsic biological meaning, but LSC1 encodes the largest distances in the matrix, LSC2 encodes the next largest distances in the matrix and so on.

### 3.4 Explantory Components

Explantory components (EC) are used to make a biological interpretation of the sample point cloud, and are graphically summarised in **Figure 4**. In more detail, a simple univariate quantity *U* is first extracted from each sample. Examples of *U* include the percentage of cells positive for a marker, average expression level of a marker amongst positive cells, and enrichment of double positive cells for marker pairs. Next, partial least squares (PLS) [4] is used to see whether the sample point cloud is linearly predictive of *U*. The information contained in an EC includes the percentage of *U* ‘s variance that is explainable by the sample point cloud, and the linear direction in the sample point cloud that maximally explains *U*. In summary, if *U* is a strong EC, then *U* will appear as a gradient across the sample point cloud.

**Figure 4:**
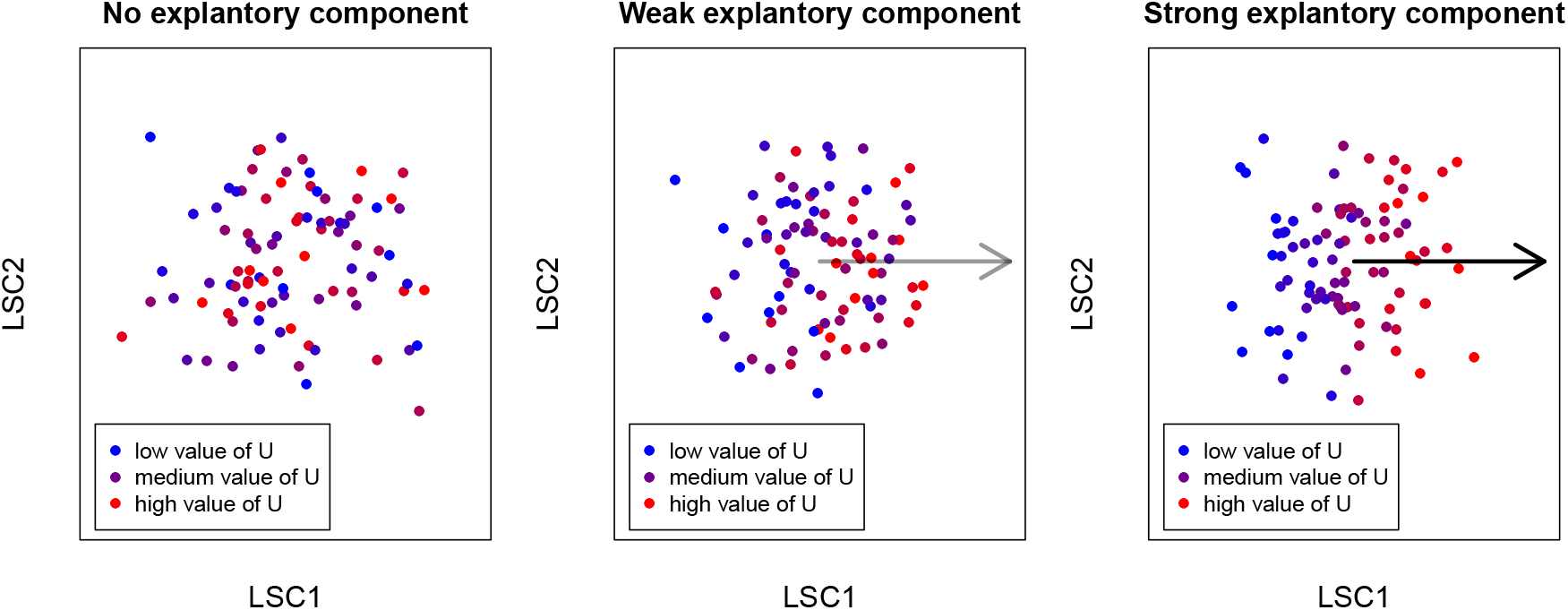
A diagram to explain the concept of explanatory components (EC). Density-Morph captures differences in complex multivariate distributions, which could be driven by intricate patterns of expression and frequency shifts over multiple markers. In order to gain an intuitive understanding of the processes that may be driving the differences, highly simplified univariate quantities *U* are extracted from each sample, and partial least squares (PLS) is used to see whether the point cloud structure is linearly predictive of *U*. The information in an EC includes the percentage of *U* ‘s variance that is explainable by the point cloud, and the linear direction in the point cloud that explains *U*. This can be understood by plotting the point cloud colour coded by the value of *U*. **Left** If the point cloud does not explain any of *U* ‘s variance, the values of *U* appear to be randomly shuffled within the point cloud. **Middle** If the point cloud explains some of *U* ‘s variance, the values of *U* will appear as a noisy gradient. **Right** If the point cloud explains most of *U* ‘s variance, the values of *U* will appear as a strong gradient. In the two latter cases, the EC is represented by an arrow aligning with the gradient and the darkness of the arrow is proportional to the percentage of *U* ‘s variance that is explained. The advantage of representing an EC with an arrow is that multiple ECs can be displayed on a single plot.

### 3.5 Trade-offs between bulk and single cell measurements

It can be helpful to think of *U* as a bulk measurement, i.e. an average measurement of a sample, in the same way a spot on a microarray is an average measurement of each gene’s expression in a sample. However, each sample has a complex multivariate structure that cannot be collapsed into a set of *U* s without a significant loss of information. Within each sample, each marker will have a complex distribution of expression values. There will also be a complex correlation structure amongst markers, implying that an expression level of 10 units for marker ‘X’ won’t have a fixed biological implication for all single cells, but that implication will vary according to the co-expressed markers. This loss of information explains why it is unlikely that any *U* can be a perfect EC for a sample point cloud. An analogy to this is that you don’t recognise a face by only the nose or eyes alone, but all the facial features work synergistically together to make a face unique.

The information gained by performing single cell experiments comes at the expense of human intuition, as we are not naturally equipped to think beyond three or four dimensions, let alone over multiple point clouds concurrently, or complex correlation structures. DensityMorph acts in a ‘black-box’ fashion by implicitly calculating over all the multivariate complexities, and the output is devoid of intuitive meaning. The meaning is the overlaid by using ECs but the user needs to be aware of the connection between *U* and the underlying multivariate structure. When interpreting the ECs, it should be kept in mind that a very strong EC is unlikely, e.g. an expression related *U* does not capture correlation with other markers but is highly influenced by it, and the correlation structure will most likely vary across the samples. On the hand, a weak EC doesn’t mean that nothing is happening, e.g. if multiple weak ECs are aligned in roughly the same direction, it could indicate a multivariate signal that is not fully captured by univariate quantities. Finally, it should be remembered that ECs currently only capture linear association between *U* and the sample point cloud, and it might be worthwhile to define a non-linear EC in the future.

### 3.6 Probing coexpression patterns using the log-odds ratio

In traditional flow cytometry manual gating, the double positive gates are usually chosen according to prior knowledge, e.g. CD3-CD4, otherwise there is a combinatorial explosion in the number of double positive gates as the number of markers increases. Additionally, simply inspecting raw percentages of double positive cells gives no indication as to whether they are over or under-represented with respect to the total number of cells positive for either marker of the pair. To probe all coexpression patterns in a cohesive manner, log-odds ratios (LOR) are used (**Figure 5**).

**Figure 5:**
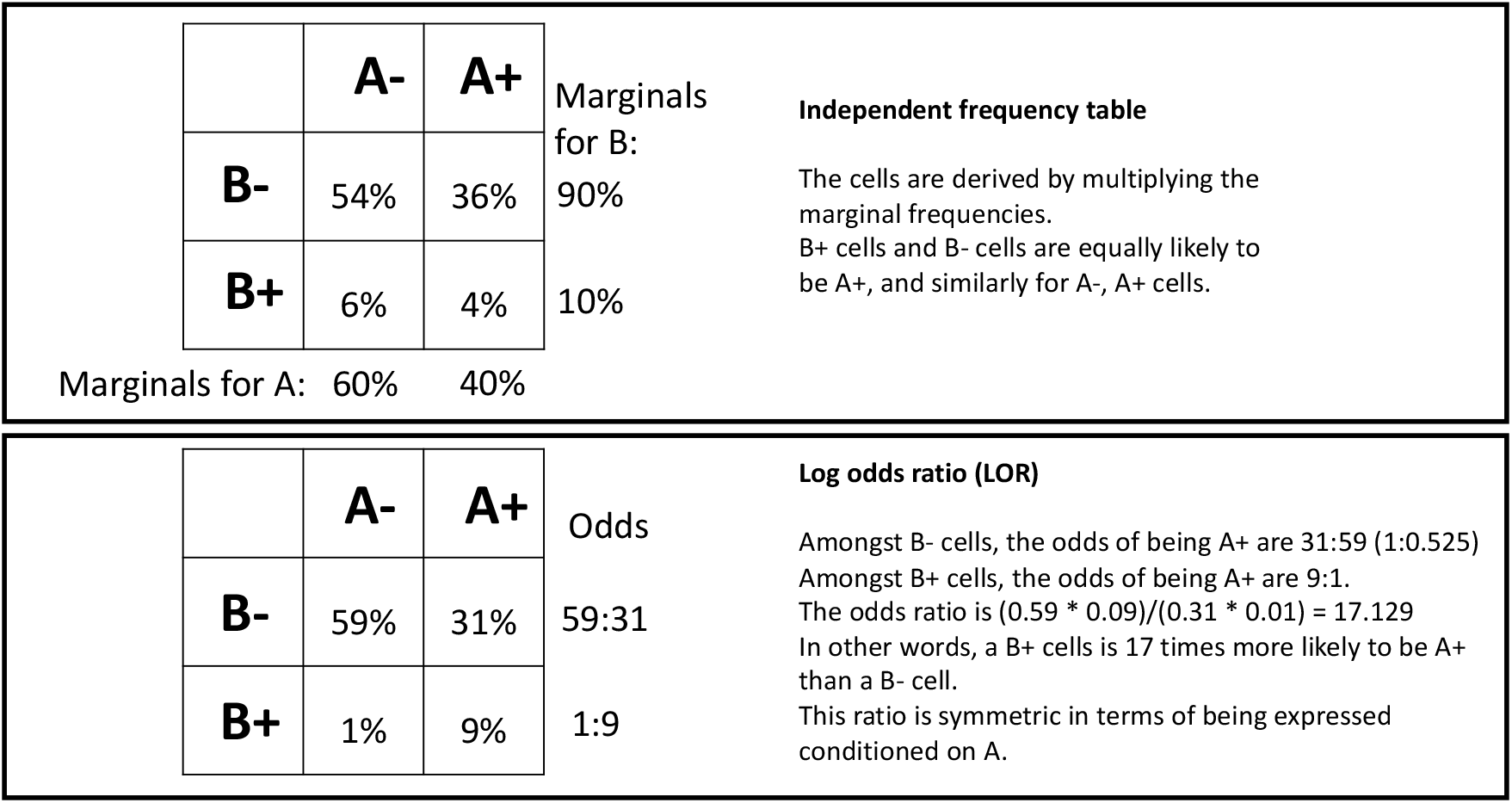
A diagram demonstrating independence and log-odds ratios (LOR) in 2 *×* 2 frequency tables. Frequency tables can summarise boolean gating results and are useful for normalising double positive frequencies relative to the underlying marginal frequencies. **Above** Markers A and B are *statistically* independent of each other, i.e. all possible combinations are obtained by multiplying the marginal frequencies. Note that ‘independence’ often has a different interpretation in statistics and biology, e.g. biological independence could be interpreted as A and B never being coexpressed on the same cell. **Below** The LOR captures the enrichment or depletion of double positive cells relative to the marginal frequencies. If the LOR is positive, there is an enrichment of AB coexpression, if the LOR is negative, there is a depletion of AB coexpression.

The LOR is derived via a 2 *×* 2 frequency table containing the percentage of single cells that are positive for marker A but not B (A^+^B^*−*^), positive for B but not A (A^*−*^B^+^), positive for A and B (A^+^B^+^), negative for A and B (A^*−*^B^*−*^). The marginal frequencies are the total percentage of A^+^ cells (whether B^+^ or B^*−*^), and similarly for B. The double positive frequency should be normalised relative to the marginal frequencies in order to uncover whether double positive cells are over or under-represented. When A and B are **statistically** independent of each other, the double positive frequency is simply the multiplication of the marginal frequencies. In this case, A^+^ and A^*−*^ cells have the same probability of also being B^+^ and the LOR is 1, and vice versa. If the LOR*<* 1, then a A^*−*^ cells is more likely to be B^+^ than a A^+^ cell; and if LOR*>* 1, then a A^*−*^ cells is less likely to be B^+^ than a A^+^ cell. The definition is symmetrical for B. Note that ‘independence’ often has a different interpretation in statistics and biology, e.g. biological independence could be interpreted as A and B never being coexpressed on the same cell.

The use of LORs as ECs allows us to see how coexpression patterns systematically vary over the samples and might point to higher order coexpression patterns. For example, if three ECs corresponding to the pairs (AB, AC, BC) are aligned in the same direction, it suggests the markers A, B and C have a strong tendency to be expressed on the same cell. It could be due to the existence of a strongly defined discrete cell population that is discoverable by a clustering algorithm. Otherwise, it could be due to a fuzzy set of biological rules where a certain cell type is positive to one, two or three of the markers, and this may not be discoverable by standard clustering algorithms. In either case, LORs are a useful tool to guide the discovery of coexpression patterns and cell populations.

## 4 Analysis of COVID data

### 4.1 Background

A dataset of COVID-19 patient flow cytometry samples is analysed [5], referred to as Lucas (2020). The dataset consists of samples taken from 113 COVID-19 patients at *≥* 1 timepoints, along with those from a number of healthcare workers, serving as healthy controls. The samples were stained with three different panels, but here I will focus on the leukocyte panel consisting of CD3, CD14, CD16, CD19, CD56, CD141, CD304, CD66b, CD1c, CD11b, CD11c, and HLA. The manual gating strategy is shown Extended Data Figure 9 of Lucas (2020), and as this is an immunophenotyping panel, the gating is straight forward with well-defined cell populations. The gating strategy is defined by well established knowledge about the major cell lineages, e.g. CD3, CD14, CD19 should all be mutually exclusive to each other as they respectively correspond to T cells, monocytes and B cells. Due to the number of markers, a exhaustive search between all marker pairs to uncover something ‘interesting’ is generally not performed when manually gating flow cytometry samples. Even a constrained gating with a known strategy is very labour intensive when performed over hundreds of samples, and further downstream work is then required to gain biological insight. Hence this is an ideal dataset to demonstrate the power of the DensityMorph method.

### 4.2 Latent space plots and experimental design

The datasets are gated to only include singlets and live cells. They are compensated and logicle-transformed. Next, dimension is reduced using Spectral Map Analysis (SMA) [6], which involves row normalisation as well as column normalisation on the log-transformed matrix, before performing PCA. This ensures that the PCA is driven by the difference in the profiles of cell populations, rather than the dynamic range of the markers. A sample-sample distance matrix is calculated using DensityMorph, the matrix is transformed element wise to the power of 0.5, and a latent space is created with PCO. The samples are plotted corresponding to their coordinates along Latent Space Components (LSC) 1 and 2 in **Figure 6**. One large cluster is evident, as well as a number of outlying samples. Reasons for outlying samples could be biological, e.g. these patients could have a common pre-existing medical condition, or technical, e.g. failure in sample preparation. The point cloud is plotted four times, with a colour code representing four selected covariates: patient vs. healthy control, gender, race and age. There is a very clear divide between patients and healthy controls. Judging by eye, there appears to be a trend related to gender and age, but not to race. However, there is a strong confounding amongst the covariates, e.g. the healthy controls are often female, young and white.

**Figure 6:**
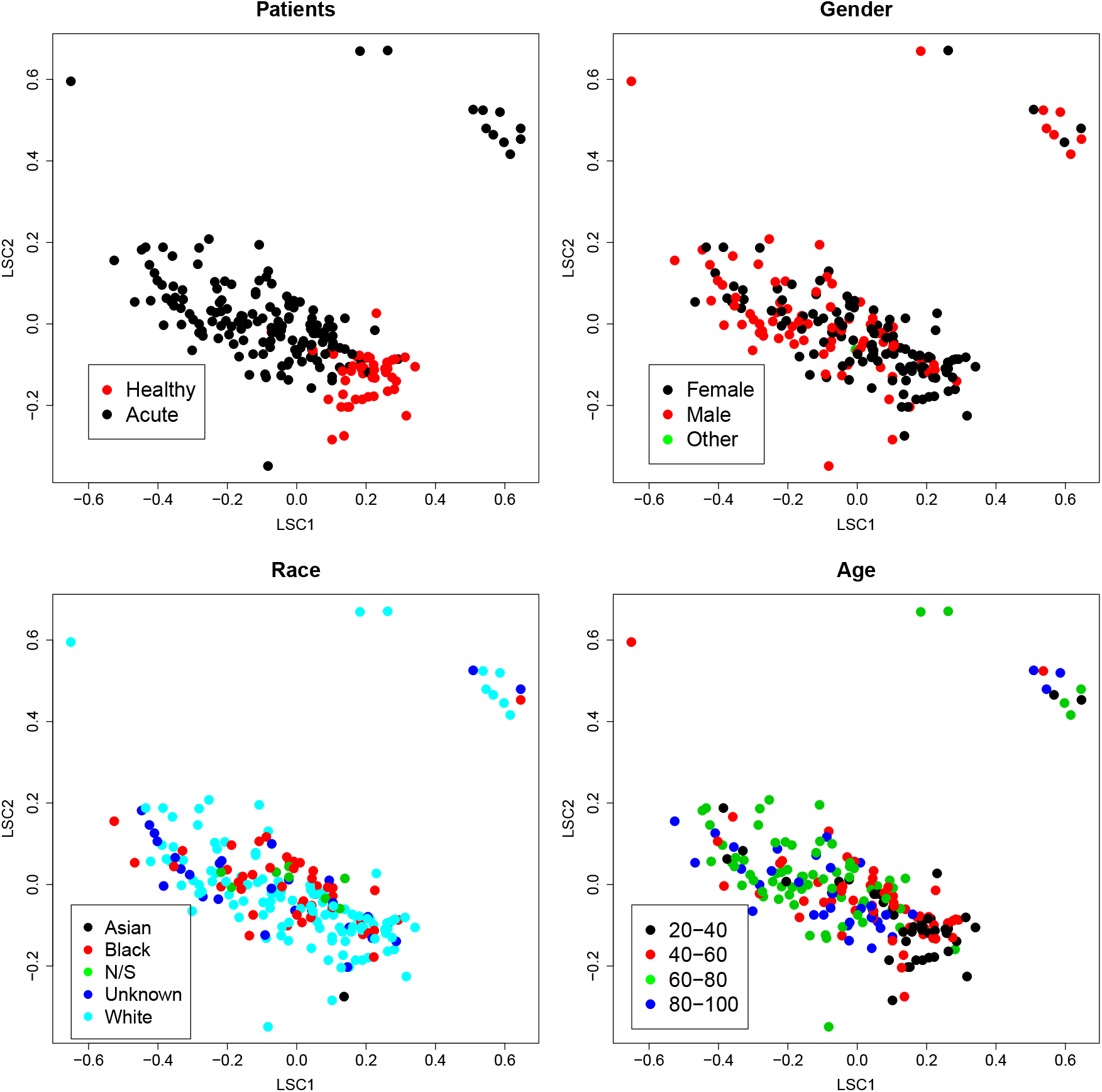
A COVID dataset consisting of 207 flow cytometry samples is embedded in a latent space by using DensityMorph. The resulting point cloud is plotted and one large cluster is evident, as well as a number of outlying samples. Reasons for outlying samples could be biological, e.g. these patients could have a common pre-existing medical condition, or technical, e.g. failure in sample preparation. The point cloud is plotted four times, with a colour code representing four selected covariates: patient vs. healthy control, gender, race and age. There is a very clear divide between patients and healthy controls. Judging by eye, there appears to be a trend related to gender and age, but not to race. However, there is a strong confounding amongst the covariates, e.g. the healthy controls are often female, young and white.

For brevity, the outlying samples are excluded in order to focus in detail on the structure of the large cluster (**Figure 6**), however the reason for being outliers should be investigated in a more thorough analysis. The latent space is re-calculated on the large cluster. There doesn’t appear to be any further structure, e.g. no new clusters have appeared, however there is a small number of mildly outlying samples in LSC2. The point cloud is again plotted four times with the four selected covariates in **Figure 7**. Information about the time courses can be found in **Figure 8** by using arrows to connect samples arising from the same patient. Many samples, including all healthy controls, are only taken at one timepoint (day zero), but other time courses range from 2 to 6 timepoints, taken at varying intervals. A small number of patients have shorter arrows, indicating a relatively stable immunophenotype over the time course. Others have much larger arrows and some don’t follow a smooth trajectory. Most notably, the trajectories show no sign of alignment with each other, indicating there is no typical disease progression amongst this group of patients. The metadata supplied with Lucas (2020) does not include information about the disease outcome so this cannot be compared with the trajectories. Finally, it should be remembered that these trajectories are only displayed in LSC1 and LSC2, and could have a substantial component in other LSCs.

**Figure 7:**
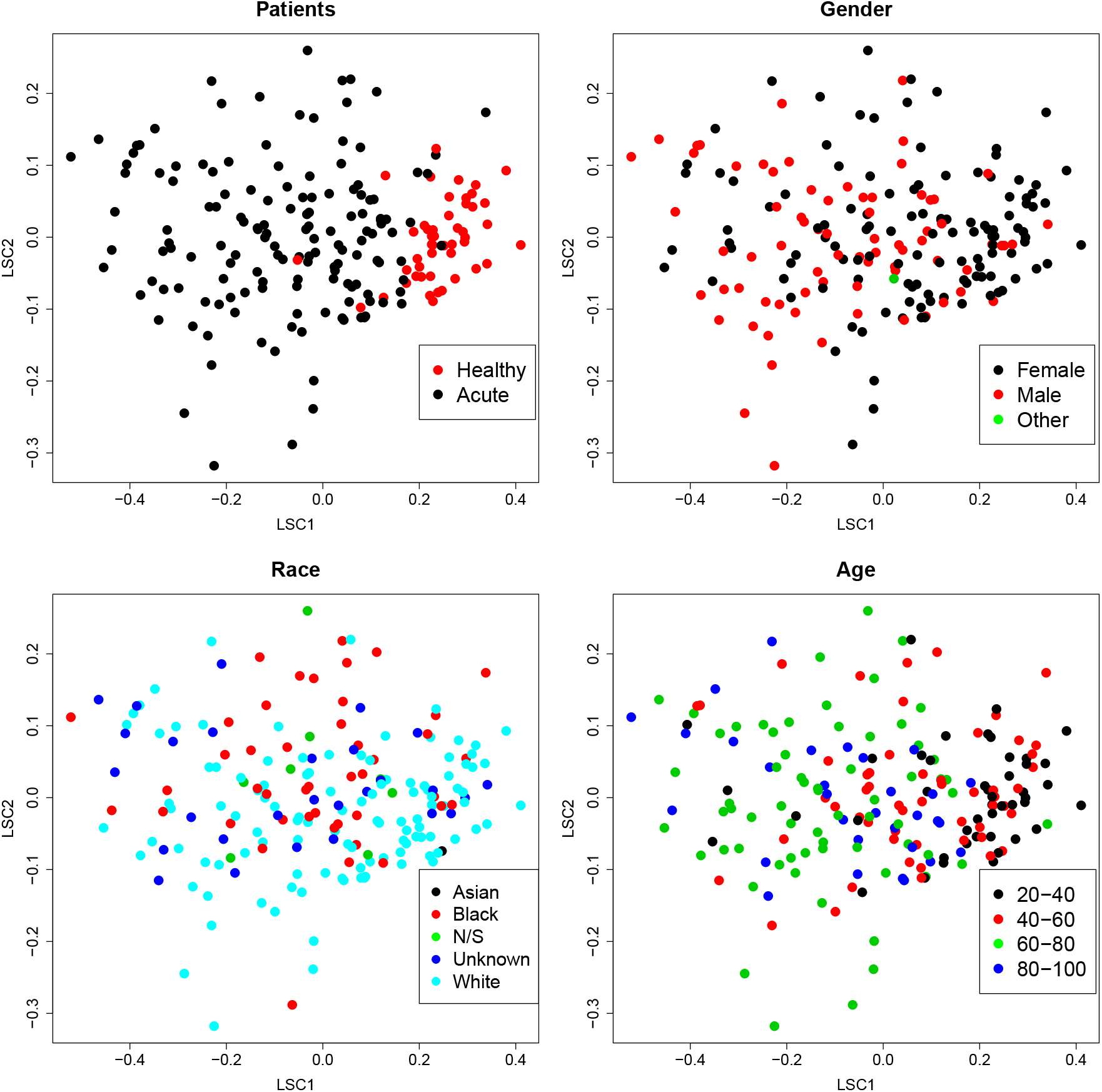
The outlying samples are excluded in order to focus in detail on the structure of the large cluster (Figure 6). The latent space is re-calculated on the large cluster. There doesn’t appear to be any further structure, e.g. no new clusters have appeared, however there is a small number of mildly outlying samples LSC2. The point cloud is again plotted four times with the four selected covariates.

**Figure 8:**
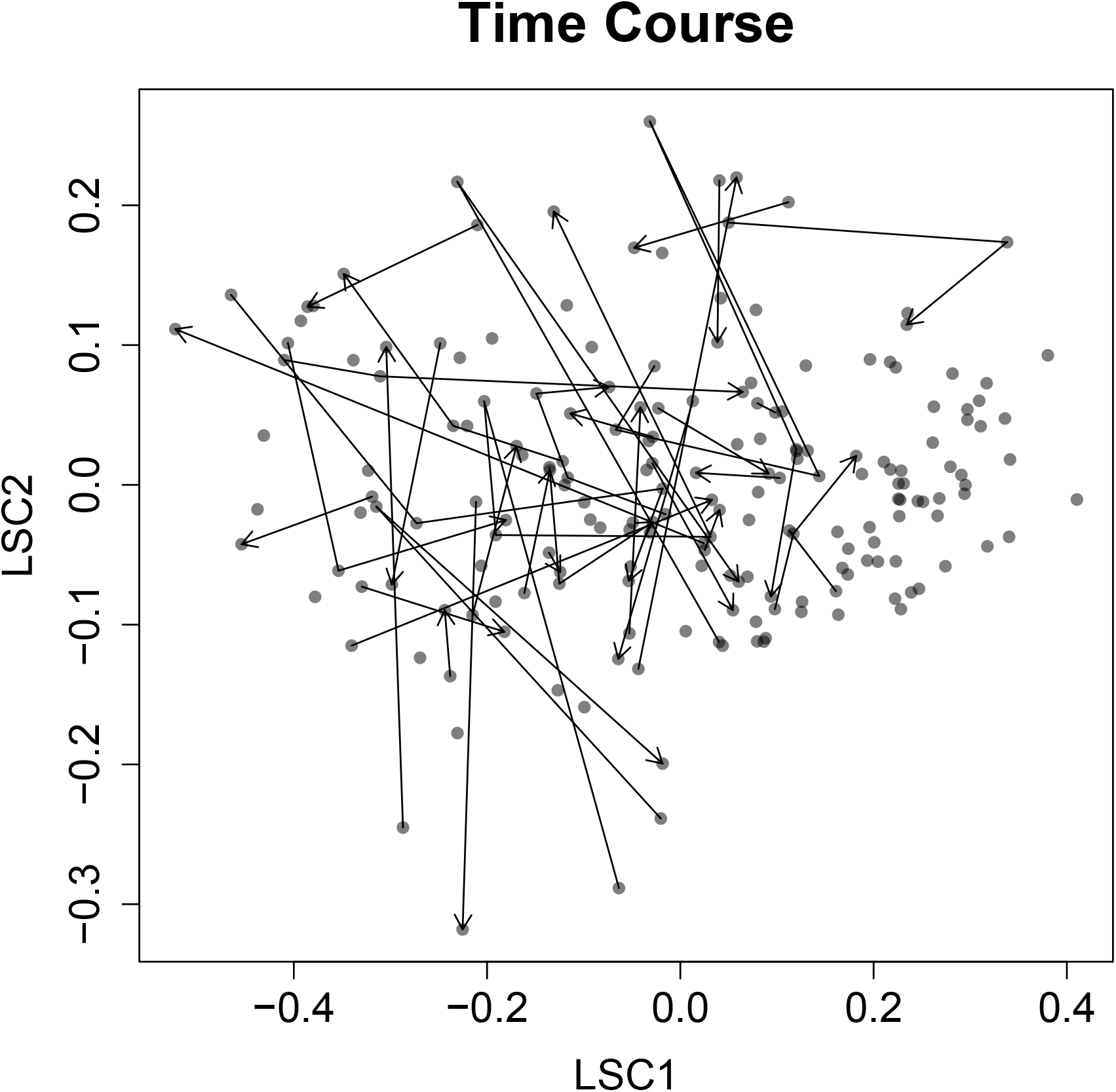
The samples are plotted in the latent space of Figure 7, with arrows representing the time courses. Many samples are only measured at one timepoint, including all the healthy controls. 25 patients are measured at 2 timepoints, 12 at 3 timepoints, 3 at 4 timepoints and one each at 5 and 6 timepoints. The time courses are taken varying intervals, and the total length of the timecourse ranges from 2 days to 26 days (out of *≥* 2 timepoints). Some time courses may encompass the excluded points shown in Figure 6 but are not shown here. The final timepoint occurs at the arrow head.

### 4.3 Expression based Explanatory Components

The first step in the interpretation of **Figure 7** is to overlay it with expression based ECs. For each sample and marker, two quantities are extracted: the percentage of positive cells (*U*_1_) and the mean expression of the positive cells (*U*_2_). The mean fluorescence intensity (MFI) of the cells is used as a proxy for the expression levels, but it in no way measures the true expression level, nor are MFI values comparable between markers. *U*_1_ and *U*_2_ can be seen as related to bulk expression measurements.

The ECs corresponding to *U*_1_ is displayed in **Figure 9**. The first LSC is driven by two anti-correlated EC bundles: (CD3, CD56), and (CD1c, CD304, CD16, CD11c, CD11b, CD66b, CD14). Referring to **Figure 7**, these ECs are aligned with patient status and age. In more detail, young healthy controls have a higher proportion of CD3^+^ and CD56^+^ cells than old sick patients, and vice versa for the second marker set. There appears to be two more gradients associated other LSCs. These correspond to HLA, and (CD141, CD19). As the point cloud is likely to be multidimensional, it is possible that some ECs may not have a strong projection in the LSC1-LSC2 plane, and this is represented by the length of the arrow. Here, the HLA EC has the shortest arrow as its two largest orthogonal projections are in LSC4 and LSC5 (**Table 1**). Referring to Extended Data Figure 2 (EDF2) of Lucas (2020), young healthy controls have significantly more T cells (count per volume), and this is in agreement with the CD3 *U*_1_-EC of **Figure 7**. The three types of dendritic cells are also more abundant in young healthy controls, but these are very rare cells and they are vastly outnumbered by T cells. As a consequence, the relative proportion of dendritic cells decreases in young healthy controls, as reflected in the *U*_1_-ECs of CD304, CD141 and CD1c. In EDF2, B cells (CD19), conventional monocytes (CD14), NK cells (CD56) and NKT cells (CD3-CD56) have either no or little significant difference in abundance between healthy and patient, however their *U*_1_- ECs show strong differences in proportion that drive the contrast between healthy and patient. Although it hasn’t been done here, there is no reason why abundance couldn’t also be used as an EC.

**Figure 9:**
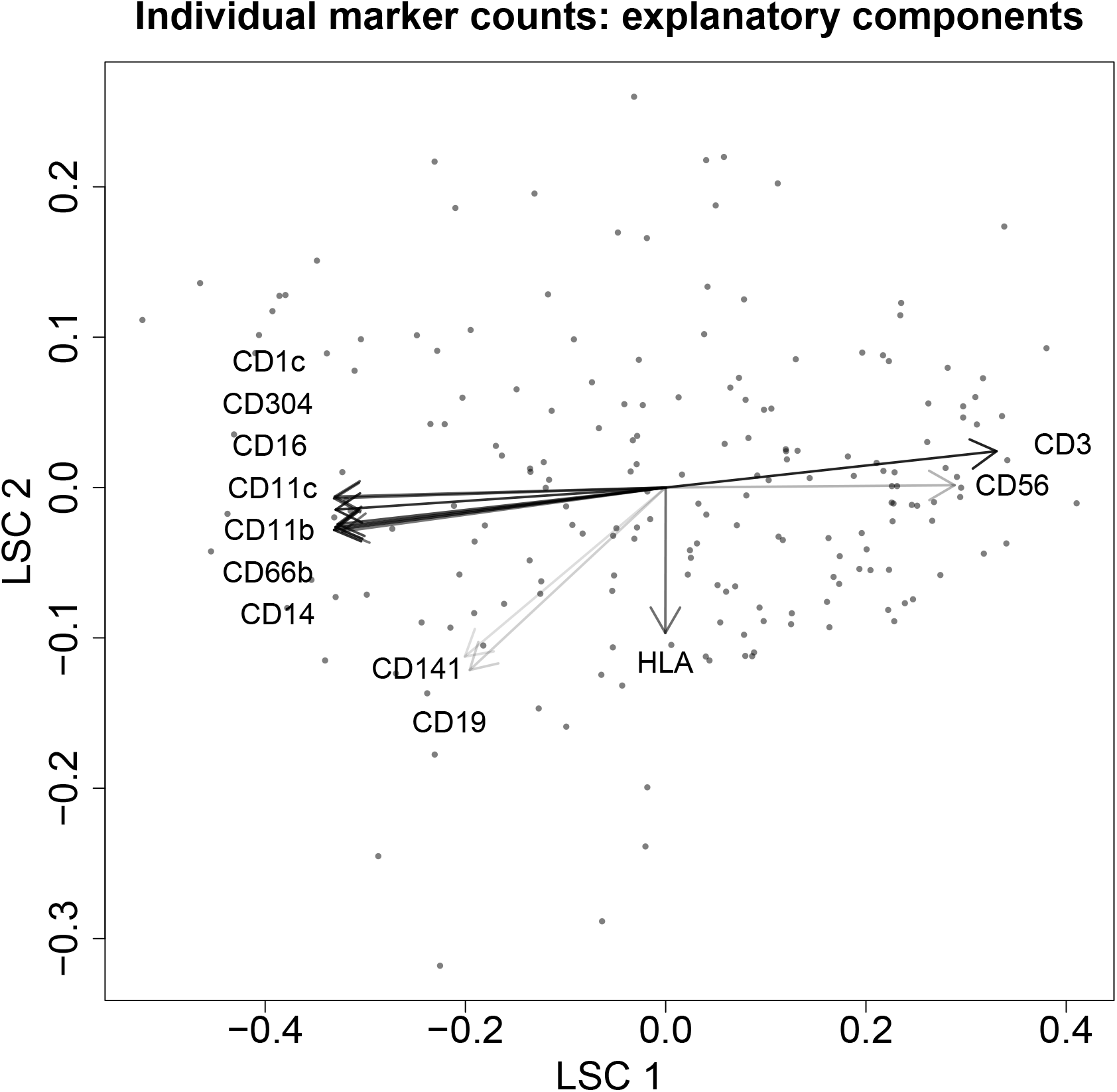
For each marker, *U*_1_ is defined to be the proportion of positive cells, regardless of the positive cells’ expression levels. The first LSC is driven by two anti-correlated EC bundles: (CD3, CD56), and (CD1c, CD304, CD16, CD11c, CD11b, CD66b, CD14). Referring to Figure 7, these ECs are aligned with patient status and age. In more detail, young healthy controls have a higher proportion of CD3 and CD56 cells than old sick patients, and vice versa for the second marker set. There appears to be two more count gradients associated other LSCs. These correspond to HLA, and (CD141, CD19). As the point cloud is likely to be multidimensional, it is possible that some ECs may not have a strong projection in the LSC1-LSC2 plane, and this is represented by the length of the arrow. Here, the HLA EC has the shortest arrow as its largest projection is in LSC4 and LSC5 (Table 1).

**Table 1:**
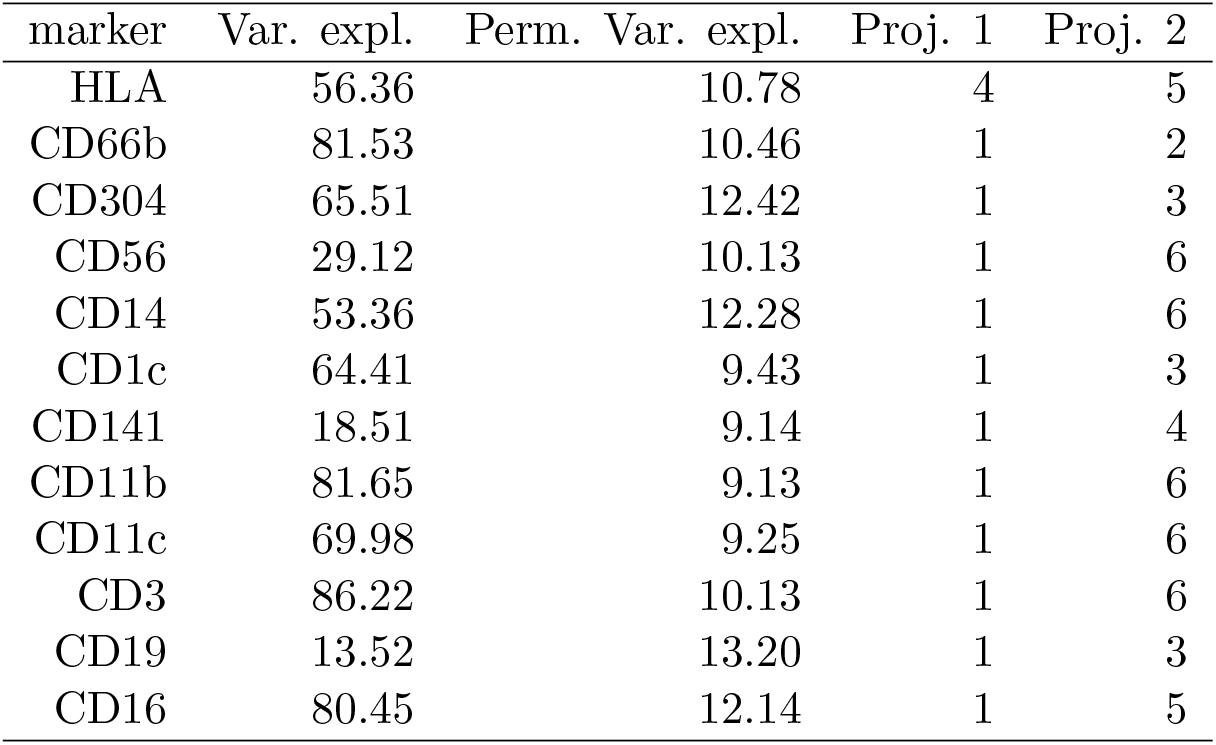
Table of *U*_1_-ECs representing the proportion of cells positive for each marker. Column 1: Marker name. Column 2: The percentage of *U*_1_’s variance explained by the point cloud. Column 3: The average percentage of the permuted *U*_1_’s variance explained by the point cloud, using 1000 permutations. Column 4: The LSC that contains the largest component of the EC. Column 5: The LSC that contains the second largest component of the EC.

The ECs corresponding to *U*_2_ is displayed in **Figure 10**. There is a bundle of ECs that appear to be aligned with younger healthy controls (CD19, CD14, CD304, CD1c, CD66b, HLA). CD3 has the strongest projection in LSC2 (**Table 2**), which seems to be driven by the small group of mildly outlying samples. Comparing to **Figure 9**, the arrangement of the *U*_2_-ECs has completely shifted compared to the *U*_1_-ECs. For example, both CD3 and CD19 flip their positions, meaning some samples have fewer T and B cells, but the cells in those samples have stronger CD3 and CD19 expression respectively. Expression levels haven’t been examined in Lucas (2020) and so no comparison can be made. This demonstrates that while ECs are easy to understand, they cannot account for the full multivariate information in the samples (**Section 3.5**). The EC plots make it straightforward to have a cohesive and integrated view of multiple aspects of the data.

**Figure 10:**
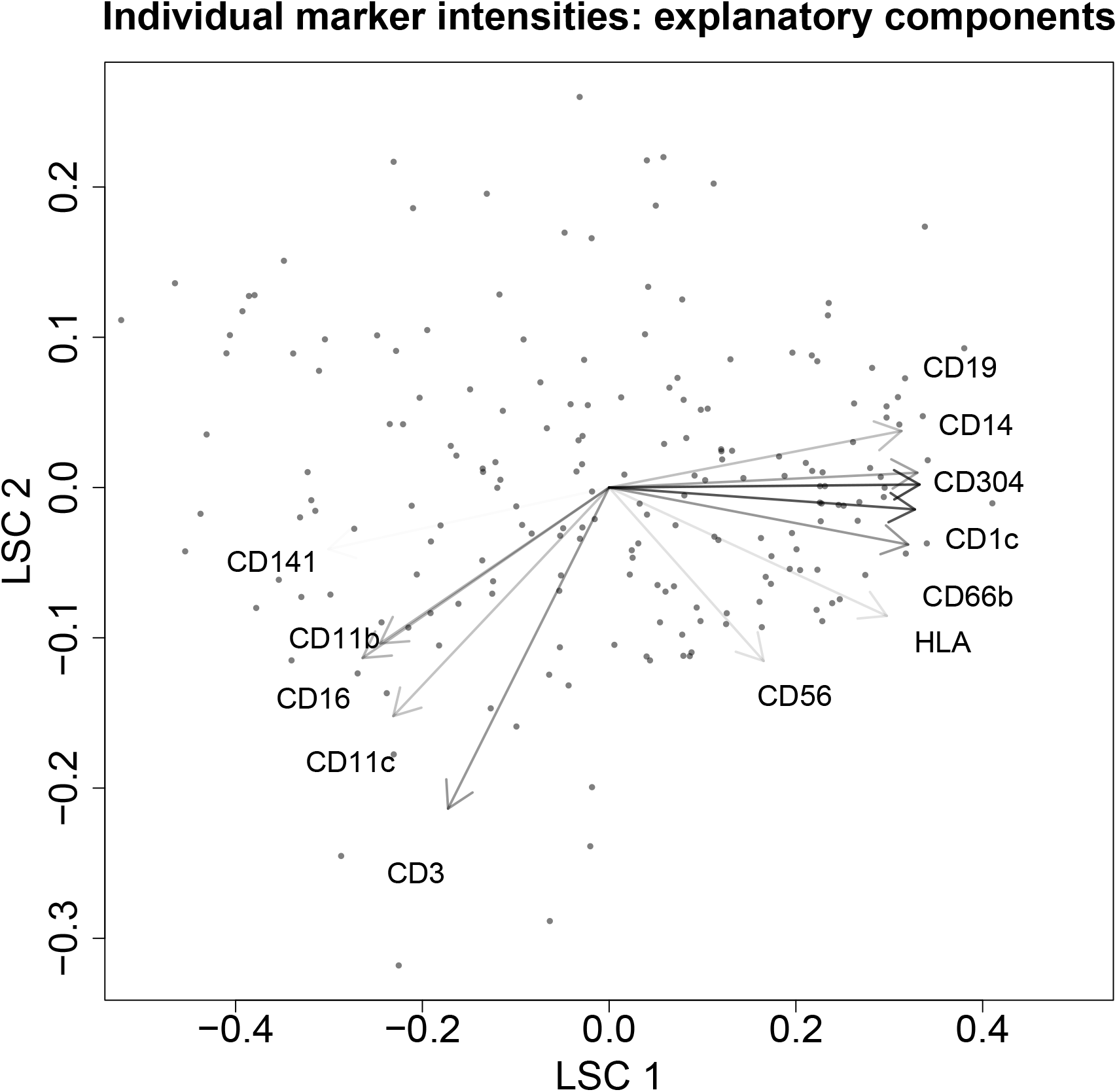
For each marker, *U*_2_ is defined to be the mean MFI of positive cells. There is a bundle of ECs that appear to be aligned with younger healthy controls (CD19, CD14, CD304, CD1c, CD66b, HLA). CD3 has the strongest projection in LSC2 (Table 2), which seems to be driven by the small group of mildly outlying samples.

**Table 2:** Table of *U*_2_-ECs representing the mean MFI of cells positive for each marker. Column 1: Marker name. Column 2: The percentage of *U*_2_’s variance explained by the point cloud. Column 3: The average percentage of the permuted *U*_2_’s variance explained by the point cloud, using 1000 permutations. Column 4: The LSC that contains the largest component of the EC. Column 5: The LSC that contains the second largest component of the EC.

| marker | Var. expl. | Perm. Var. expl. | Proj. 1 | Proj. 2 |
| --- | --- | --- | --- | --- |
| HLA | 10.90 | 10.01 | 1 | 2 |
| CD66b | 40.93 | 11.17 | 1 | 4 |
| CD304 | 35.56 | 13.09 | 1 | 7 |
| CD56 | 11.67 | 9.40 | 6 | 1 |
| CD14 | 66.75 | 12.55 | 1 | 4 |
| CD1c | 71.36 | 9.25 | 1 | 4 |
| CD141 | 1.75 | 14.18 | 1 | 9 |
| CD11b | 28.52 | 13.52 | 1 | 3 |
| CD11c | 23.69 | 12.52 | 1 | 2 |
| CD3 | 40.96 | 12.10 | 2 | 1 |
| CD19 | 24.63 | 11.15 | 1 | 3 |
| CD16 | 29.45 | 12.34 | 1 | 2 |

### 4.4 Coexpression Explanatory Components

Cell populations are rarely, if ever, defined exclusively by one single marker. The specific combination of markers expressed on or in a cell endows it with a specific biological function, and the marker combination may be fixed or transient. Furthermore, cell ‘populations’ aren’t necessarily discrete but rather cells may exist on a continuum between fixed points depending on the rules of the underlying dynamical system. The variation away from fixed populations may be induced by the temporal evolution of cell states, or it may be fixed over certain time scales and the variation itself contributes to the biological function. It is for these reasons that single cell measurements have become so prominent, as this information is obscured by bulk measurements. While the average expression ECs in **Section 4.3** were informative, for the same reasons it is necessary to also consider how coexpression patterns drive differences between samples. This will be done by using LOR scores (**Section 3.6**) as ECs. They will be used over all possible marker pairs without input from prior knowledge about known cell populations.

Firstly, LOR scores are calculated for all marker pairs over all samples and presented as a boxplot in **Figure 11**. Marker pairs that are generally considered to be mutually exclusive such as CD3-CD14 (T cells, monocytes) or CD3-CD19 (T cells, B cells) reassuringly have LOR scores that are overwhelmingly negative. As the positive/negative threshold for each marker hasn’t been set individually for each sample, there are occasions where these marker pairs will have positive LORs as the threshold may not be precise enough. Marker pairs with an interquartile range (IQR) being negative are omitted as potential ECs to reduce the number to consider, although it’s possible that some contain interesting trends.

**Figure 11:**
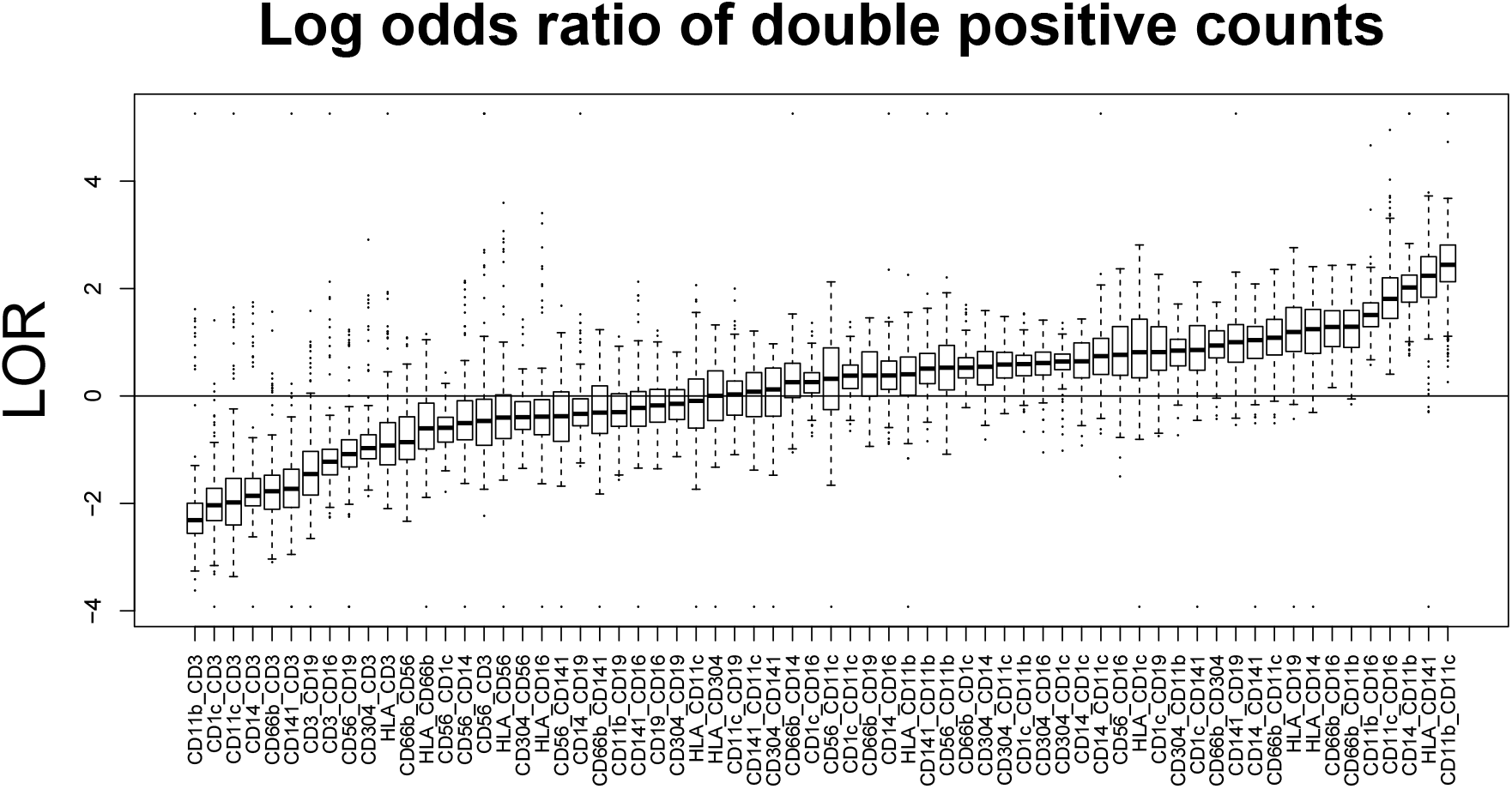
LOR scores are calculated for all marker pairs over all samples and presented as a boxplot. Marker pairs that are generally considered to be mutually exclusive such as CD3-CD14 (T cells, monocytes) or CD3-CD19 (T cells, B cells) reassuringly have LOR scores that are overwhelmingly negative. As the positive/negative threshold for each marker hasn’t been set individually for each sample, there are occasions where these marker pairs will have positive LORs. Marker pairs with an interquartile range (IQR) being negative are omitted as potential ECs to reduce the number to consider, although it’s possible that some contain interesting trends.

For each marker, *U*_3_ is defined to be the LOR of marker pairs defined in Figure 11, and the corresponding ECs are plotted in **Figure 12**, and results are in **Tables 3 and 4**. While the boxplot (**Figure 11**) was used to omit some marker pairs without knowledge about cell populations, there is a chance that some meaningless combinations have remained. Checking the tables, the *U*_3_-EC for CD14-CD19 doesn’t explain much more variance that the permuted EC, and this is because the marker pair should be mutually exclusive (monocytes and B cells). The *U*_3_-EC for CD304-CD1c explains less variance than the permuted EC, and these markers correspond to pDC and cDC2 cells.

**Figure 12:**
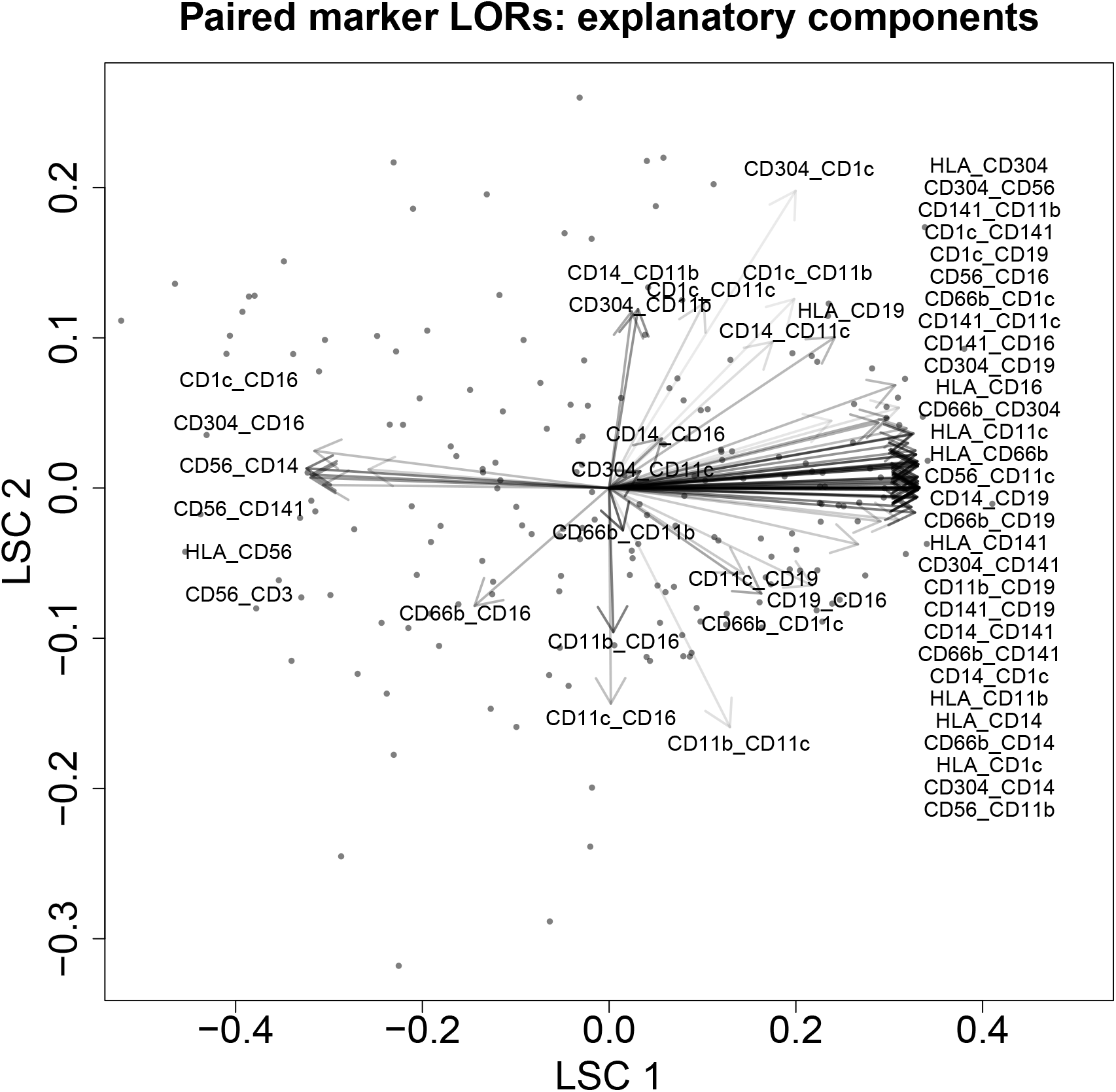
For each marker, *U*_3_ is defined to be the LOR of marker pairs defined in Figure 11. The first LSC is driven by two anti-correlated EC bundles, where the majority of the ECs are aligned with young healthy controls. An EC corresponding to NKT cells (CD3-CD56) is aligned with patients. The second LSC is driven a smaller number of more diffuse ECs. For LSC2*<* 0 the coexpression patterns involve CD16 and CD66 (neutrophils), and for LSC2*>* 0, they involve CD14, CD304 and CD1c (dendritic cells).

**Table 3:** Table of *U*_3_-ECs representing the LOR score for each marker pair (Part 1). Column 1: Marker names. Column 2: The percentage of *U*_3_’s variance explained by the point cloud. Column 3: The average percentage of the permuted *U*_3_’s variance explained by the point cloud, using 1000 permutations. Column 4: The LSC that contains the largest component of the EC. Column 5: The LSC that contains the second largest component of the EC.

| marker | Var. expl. | Perm. Var. expl. | Proj. 1 | Proj. 2 |
| --- | --- | --- | --- | --- |
| HLA_CD66b | 64.33 | 10.81 | 1 | 4 |
| HLA_CD304 | 67.78 | 12.78 | 1 | 6 |
| HLA_CD56 | 50.67 | 9.55 | 1 | 5 |
| HLA_CD14 | 50.85 | 14.58 | 1 | 9 |
| HLA_CD1c | 83.53 | 10.75 | 1 | 4 |
| HLA_CD141 | 19.48 | 10.32 | 1 | 4 |
| HLA_CD11b | 53.66 | 9.52 | 1 | 5 |
| HLA_CD11c | 41.05 | 11.50 | 1 | 6 |
| HLA_CD19 | 27.76 | 13.64 | 1 | 3 |
| HLA_CD16 | 55.64 | 12.50 | 1 | 10 |
| CD66b_CD304 | 41.79 | 10.62 | 1 | 6 |
| CD66b_CD14 | 33.08 | 10.27 | 1 | 4 |
| CD66b_CD1c | 27.47 | 13.82 | 1 | 6 |
| CD66b_CD141 | 70.93 | 12.96 | 1 | 4 |
| CD66b_CD11b | 60.17 | 9.27 | 4 | 3 |
| CD66b_CD11c | 35.03 | 10.31 | 3 | 1 |
| CD66b_CD19 | 50.05 | 10.02 | 1 | 3 |
| CD66b_CD16 | 29.29 | 11.16 | 3 | 1 |
| CD304_CD56 | 58.29 | 10.39 | 1 | 5 |
| CD304_CD14 | 55.34 | 9.42 | 1 | 4 |
| CD304_CD1c | 8.37 | 9.80 | 1 | 2 |
| CD304_CD141 | 63.78 | 9.51 | 1 | 6 |
| CD304_CD11b | 43.06 | 11.63 | 5 | 3 |
| CD304_CD11c | 53.21 | 9.07 | 3 | 4 |
| CD304_CD19 | 20.80 | 14.84 | 1 | 3 |
| CD304_CD16 | 26.43 | 12 | 1 | 6 |

**Table 4:** Table of *U*_3_-ECs representing the LOR score for each marker pair (Part 2). Column 1: Marker names. Column 2: The percentage of *U*_3_’s variance explained by the point cloud. Column 3: The average percentage of the permuted *U*_3_’s variance explained by the point cloud, using 1000 permutations. Column 4: The LSC that contains the largest component of the EC. Column 5: The LSC that contains the second largest component of the EC.

| marker | Var. expl. | Perm. Var. expl. | Proj. 1 | Proj. 2 |
| --- | --- | --- | --- | --- |
| CD56_CD14 | 18.56 | 11.37 | 1 | 5 |
| CD56_CD141 | 44.77 | 10.57 | 1 | 5 |
| CD56_CD11b | 69.33 | 13.28 | 1 | 8 |
| CD56_CD11c | 80.66 | 10.73 | 1 | 3 |
| CD56_CD3 | 40.65 | 12.17 | 1 | 7 |
| CD56_CD16 | 77.21 | 9.89 | 1 | 8 |
| CD14_CD1c | 68.64 | 10.98 | 1 | 4 |
| CD14_CD141 | 12.65 | 10.37 | 1 | 9 |
| CD14_CD11b | 33.98 | 13.83 | 5 | 2 |
| CD14_CD11c | 13.37 | 10.32 | 1 | 7 |
| CD14_CD19 | 9.71 | 7.72 | 1 | 6 |
| CD14_CD16 | 36.26 | 10.96 | 7 | 6 |
| CD1c_CD141 | 85.67 | 12.29 | 1 | 4 |
| CD1c_CD11b | 8.60 | 9.65 | 1 | 2 |
| CD1c_CD11c | 16.95 | 10.81 | 4 | 5 |
| CD1c_CD19 | 68.36 | 10.76 | 1 | 4 |
| CD1c_CD16 | 25.89 | 10.96 | 1 | 5 |
| CD141_CD11b | 26.55 | 10.58 | 1 | 5 |
| CD141_CD11c | 38.10 | 11 | 1 | 2 |
| CD141_CD19 | 31.46 | 11.64 | 1 | 3 |
| CD141_CD16 | 50.52 | 9.13 | 1 | 5 |
| CD11b_CD11c | 12.15 | 9.89 | 6 | 2 |
| CD11b_CD19 | 21.67 | 9.92 | 1 | 3 |
| CD11b_CD16 | 49.61 | 11.63 | 4 | 6 |
| CD11c_CD19 | 22.04 | 11.47 | 3 | 4 |
| CD11c_CD16 | 24.71 | 10.31 | 6 | 2 |
| CD19_CD16 | 18.33 | 10.57 | 1 | 3 |

The first LSC is driven by two anti-correlated EC bundles. The majority of *U*_3_-ECs are aligned with young healthy controls, indicating that patients’ cells have lost the ability to co-express markers relative to young healthy controls. The oldest patients are aligned with a *U*_3_-EC for CD3-CD56, which correspond to NKT cells. Interestingly, this EC is anti-correlated to the *U*_1_-ECs for CD3 and CD56 (**Figure 9**). This means that while patients have a lower proportion of CD3^+^ and CD56^+^ cells overall, when these two markers are expressed on patients’ cells, they are more likely to be coexpressed (NKT cells) than be mutually exclusive (T cells and NK cells). However it is not possible to say from this plot alone whether patients have a higher proportion of NKT cells than healthy controls. Other *U*_3_-ECs aligned with patients show that CD56 is also preferentially coexpressed with CD1c, CD14, CD141, HLA. The second LSC is driven by some smaller and more diffuse bundles of *U*_3_-ECs. For LSC2*<* 0 the coexpression patterns involve CD16 and CD66, corresponding to neutrophils, and for LSC2*>* 0, they involve CD14, CD304 and CD1c, corresponding to dendritic cells.

## 5 Summary and Outlook

I have presented an exploratory data analysis and visualisation method that can summarise a large number of single cell datasets with very minimal prior assumptions. For a first pass, it is not necessary to have prior knowledge about gating strategies or cell population structure, as it was demonstrated in the example above that this information is implicitly incorporated into the analysis. Three quantities *U*_1_, *U*_2_, and *U*_3_ are used as ECs to interpret the latent space plot but this is by no means an exhaustive list. Other quantities could also be used such as absolute abundance (as opposed to proportion), or external covariates such as plasma cytokine concentration which is also measured in Lucas (2020). The EC examples given do not necessarily lead to complete information about the dataset but it can lead to designing strategies for gating or clustering - which in turn can lead to more targeted ECs such very specific cell subsets. One example of this occurred when the *U*_1_-ECs for CD3 and CD56 were anti-correlated with the *U*_3_-EC for CD3-CD56, leading to the hypothesis that the proportions of T Cells and NK cells are anti-correlated to the proportion of NKT cells, but this would need to be confirmed by going back to the raw data.

As a consequence of the flexibility of this method, there are many future opportunities for improvement and extension. One obvious avenue is to improve the treatment of coexpression patterns. Firstly, the presented method (**Section 4.4**) works well for flow cytometry data because of the relatively low number of markers but some kind of compression scheme would need to be implemented to use this with scRNAseq data. Judging **Figure 12** by eye, it can be seen that correlated ECs have a tendency to overlap in markers. This points to a new type of single cell coexpression analysis, where each marker pair is a node on a graph, and nodes are connected if the corresponding EC is correlated above a certain threshold. The structure of the resulting graph can then be analysed using standard techniques to understand whether there are well-defined co-expression modules. This can be integrated with prior biological knowledge or used to define functionally relevant cell subsets in less well characterised cells such as cancer associated fibroblasts [7].

While this method has been presented as exploratory and visual, it could be built into a more formal inference framework, e.g. for clinical diagnostics. The number of PCO components would need to be determined in order to represent the samples in a basis of small but sufficient dimension, to allow for regression (linear or non-linear) with external covariates. The regression results would need to be linked to ECs that are calculated directly from the samples and this could be done by sparsely approximating the samples by selected ECs. This could lead to complex multivariate biomarkers that have higher precision in a clinical setting than the more usual univariate biomarkers.

I hope this manuscript has demonstrated the power of an integrated, top-down view of many single cell samples. It has the potential to amplify weak and noisy univariate signals that have been taken out of their multivariate context, such as displaying coarse-grained cell population counts with boxplots. This method can also be used to design downstream experiments - for maximum reproducibility it might be wise to experimentally compare maximally different samples, or otherwise increase the number of replicates when experimentally comparing neighbouring samples. This would be achieved to greatest effect by performing single cell samples collectively over many labs with a common interest, to map out as many of the allowable biological states as possible, before proceeding to more detailed experiments.

## 8 Supplementary Information

### 8.1 DensityMorph algorithm

Let there be two arbitrary *p*-dimensional sets of *N*_1_, *N*_2_ *p*-dimensional points 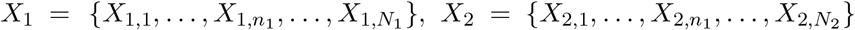, where *N*_1_ = *N*_2_. If *N*_1_ ≠ *N*_2_, then the set with the greatest number of points should be subsampled such that it contains min(*N*_1_, *N*_2_) points. Define the quasi-distance DM(*X*_1_, *X*_2_) as follows:

#### Paired density logratio

For *p*-dimensional point sets *P* and *Q* of *N* points each and *i, j ∈ {*1, …, *N}*:

1. *D*_*i*_ = min_*j*_ *‖ P*_*i*_ *− Q*_*j*_ *‖* _2_ and *D* = {*D*_1_, …, *D*_*i*_, …, *D*_*N*_}, i.e. the set of cross-NN distances from *P* to *Q*.
2. *d*_*i*_ = min_*k∈{*1,…,*N}\i ‖*_ *P*_*i*_ *− P*_*k ‖* 2_ and *d* = *{d*_1_, …, *d*_*i*_, …, *d*_*N*_}, i.e. the set of NN distances within *P*
3. PDLR_*i*_(*P*_*i*_, *Q*) = log(*d*_*i*_*/D*_*i*_) and PDLR(*P, Q*) = *{*PDLR_1_(*P*_1_, *Q*), …, PDLR_*i*_(*P*_*i*_, *Q*), …, PDLR_*i*_(*P*_*i*_, *Q*)}, i.e. the set of paired density logratios for *P* with respect to *Q*

#### DensityMorph

Let *Z*_1_, *Z*_2_ be sets of *N p*-dimensional points drawn from a *p*-dimensional normal distribution of fixed mean and variance and correlation of zero.

1. *G*_ref_ = PDLR(*Z*_1_, *Z*_2_) (self-PDLR)
2. *G*_1_ = PDLR(*X*_1_, *X*_2_)
3. *G*_2_ = PDLR(*X*_2_, *X*_1_)
4. DM(*X*_1_, *X*_2_) = max(WD(*G*_1_, *G*_ref_), WD(*G*_2_, *G*_ref_)), where WD is the 1-dimensional wasserstein distance.

The rationale for *G*_ref_ will be explained in **Section 8.3**. In practice, *G*_ref_ only needs to be calculated once for each *p*. In my implementation, I have calculated *G*_ref_ using 100*N* points and used *N* evenly spaced quantiles to have a smoother univariate distribution. In the future, *G*_ref_ may be able to be expressed in analytical form.

### 8.2 Computational Time

The computational time of the wasserstein distance (or transport distance) is investigated in **Figure 13** as a function of the number of points in each set *N* and dimension *p*. For *p ∈ {*1, 2, 3, 4, 5}, a point cloud *X*_1_ of *N* points, mean **0**, variance of 1 and correlation of 0 is compared to point cloud *X*_2_ of *N* points, mean (1, **0**), variance of 1 and correlation of 0. For *p >* 1, the ‘networkflow’ implementation is used [8]. For *N* = 10^4^, the difference in computational time between *p* = 1 and *p >* 1 is more than 4 orders of magnitude - or the difference between 1 second and 2.77 hours. This means it is impractical to use on a routine basis for the comparison of single cell samples. On that basis, I decided to use nearest neighbour (NN) distances to see if I could make an adequate approximation of the *p*-wasserstein distance. For *N* = 10^4^, the computational time is still 2-3 orders of magnitude lower (**Figure 14**), depending on *p*, and so the DensityMorph approximation will be very advantageous.

**Figure 13:**
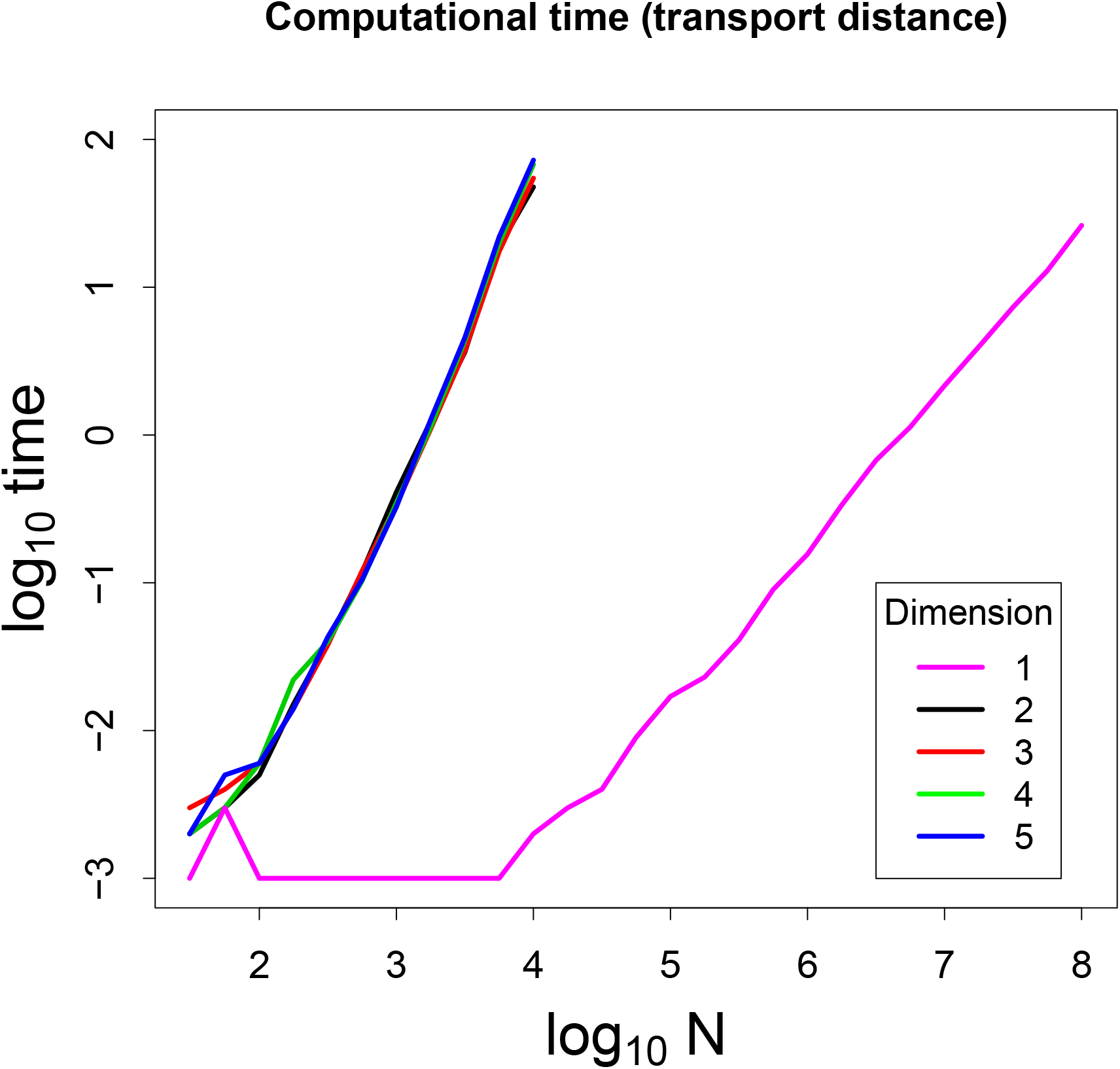
Computational time of calculating transport distance between two sets of points, as a function of number of points in each set *N* and dimension *p*.

**Figure 14:**
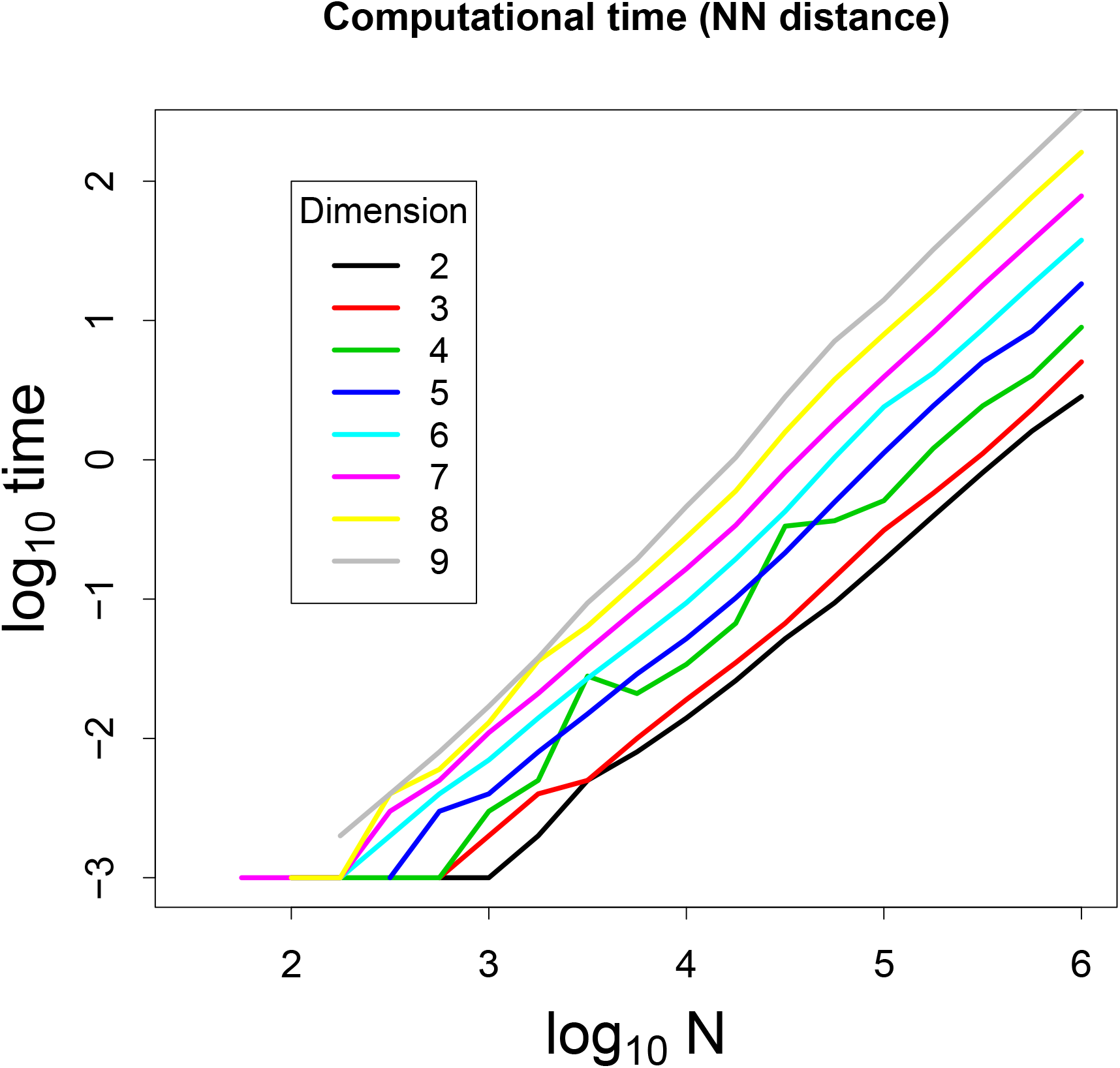
Computational time of calculating nearest neighbour (NN) distance from each point in a set to another set of points, as a function of number of points in each set *N* and dimension *p*.

### 8.3 Rationale for *G*_ref_

#### Proposition

For two sets *Y*_1_ and *Y*_2_ of *N p*-dimensional points arising from the same arbitrary mixture of gaussian distributions, the mean of PDLR(*Y*_1_, *Y*_2_) tends to zero as *N* increases, and the variance of PDLR(*Y*_1_, *Y*_2_) tends to approximately 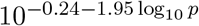 (**Figure 17**) as *N* increases.

To arrive computationally at this result, the self-PDLR distributions are calculated for paired sets of *N p*-dimensional points, arising from a *p*-dimensional normal distribution with fixed mean, correlation of zero, variance *∈ {*1, 5, 10*}, N ∈ {*100, 500, 1000, 5000, 10000}, and *p ∈ {*2, 3, 4, 5}. Each self-PDLR distribution is fit with a 1-dimensional normal distribution to estimate the mean and variance, although it is not assumed that the self-PDLR distribution is necessarily normal. This is repeated 100 times for each parameter combination and the results are displayed as boxplots in **Figures 15 and 16**. As *N* increases, the mean seems to converge on zero, and the variance seems to converge on a *p*-dependent value. The median variance is plotted against *p* in **Figure 17**. The same procedure is repeated with arbitrary mixtures of *p*-dimensional normal distributions (**Figures 18 and 19**), and again the means seem to converge to zero, and the variance seems to converge to those calculated from **Figure 17**. It suggests that the self-PDLR distribution is universal, dependent only on *p*.

**Figure 15:**
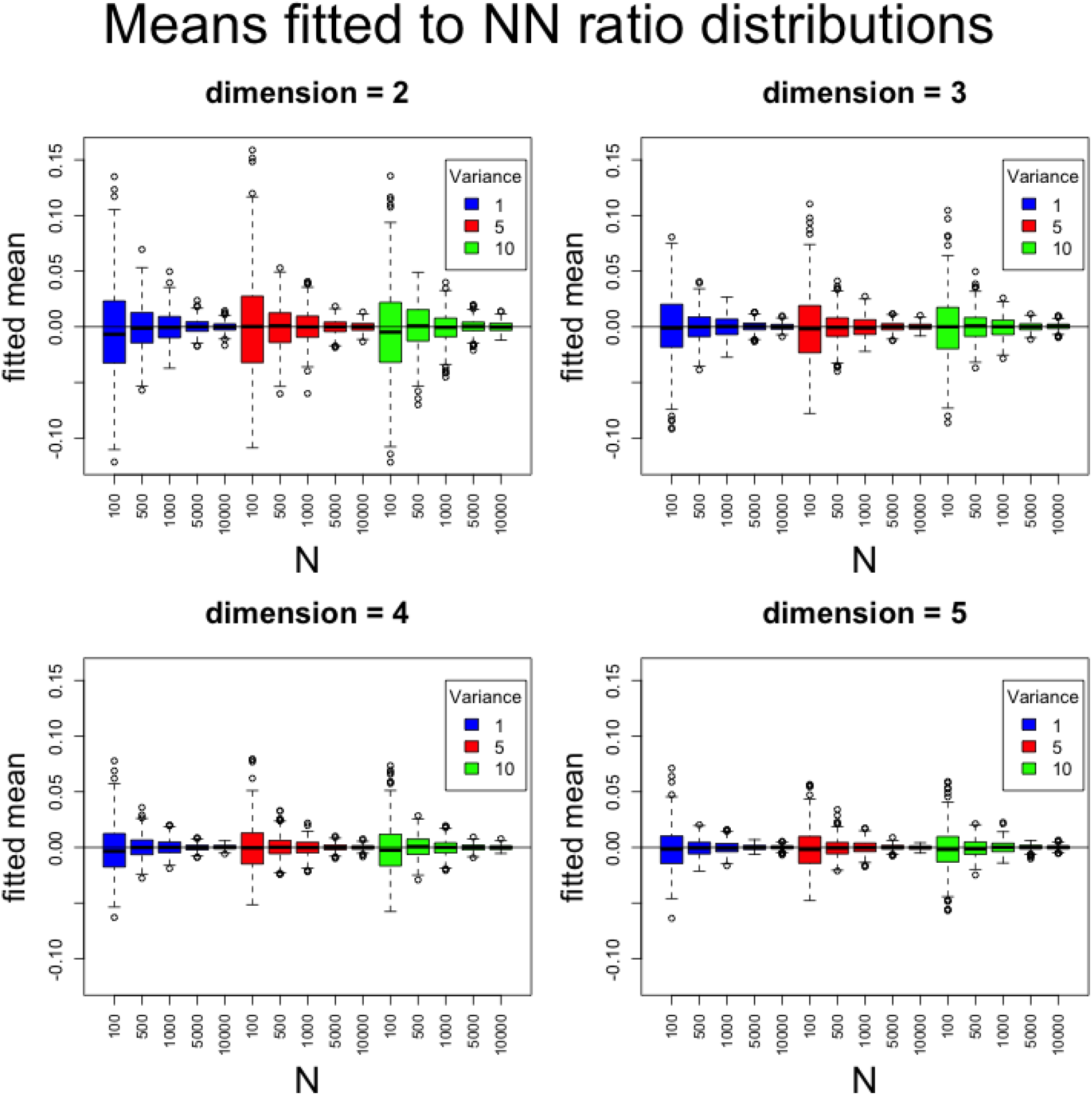
Two sets of *N* points are drawn from a *p*-dimensional normal distribution with mean of **0**, fixed variance and correlation of zero. The paired density ratio is calculated and fit using a single gaussian, and this is repeated 100 times for each parameter combination. The fitted means are displayed as a boxplot over all the parameters. The fitted means tend towards zero with increasingly smaller variance as *N* increases.

**Figure 16:**
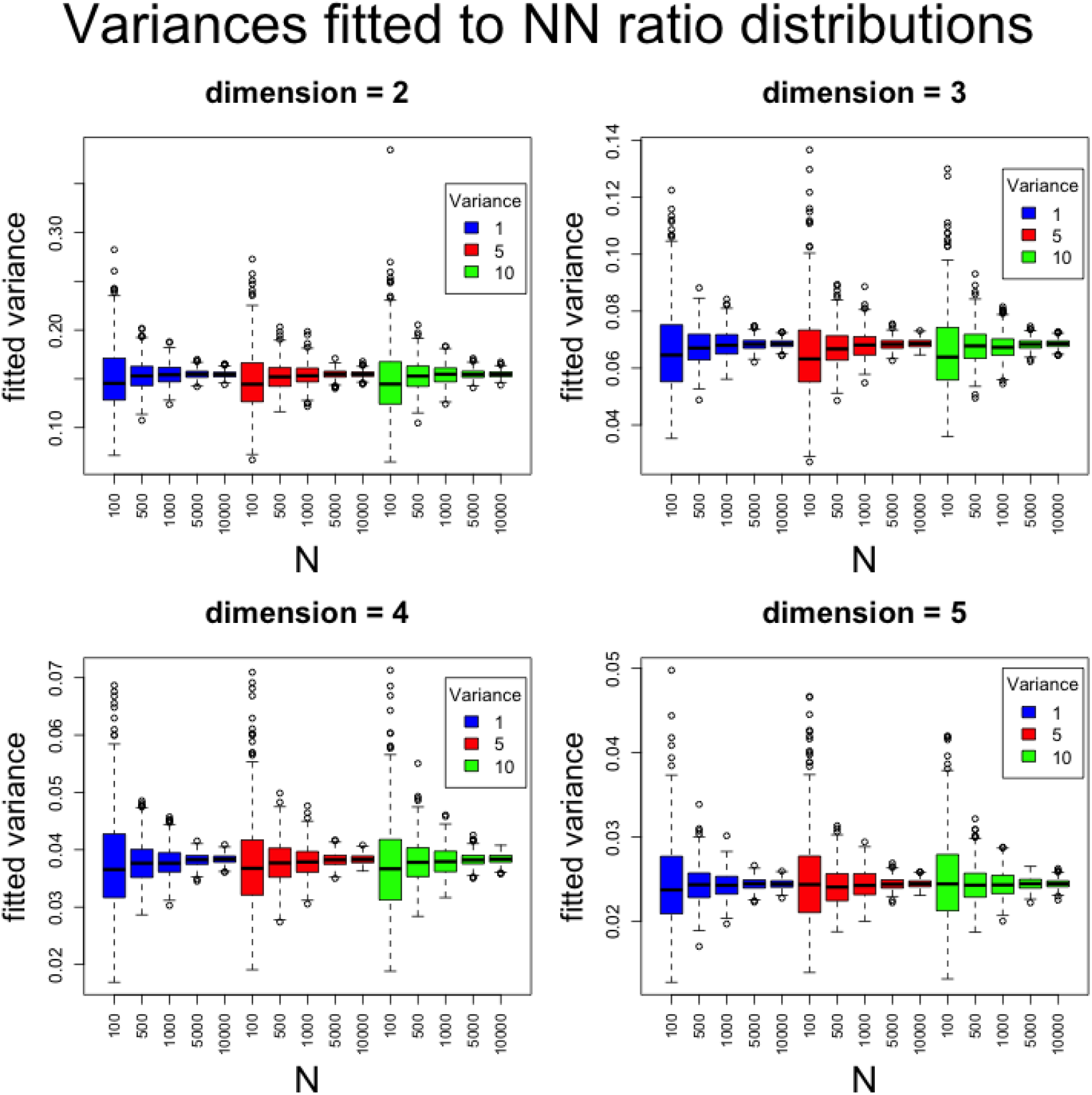
Two sets of *N* points are drawn from a *p*-dimensional normal distribution with mean of **0**, fixed variance and correlation of zero. The paired density ratio is calculated and fit using a single gaussian, and this is repeated 100 times for each parameter combination. The fitted variances are displayed as a boxplot over all the parameters. For each value of *p*, the fitted variances tend towards a fixed value with increasingly smaller variance as *N* increases. This value decreases as *p* increases.

**Figure 17:**
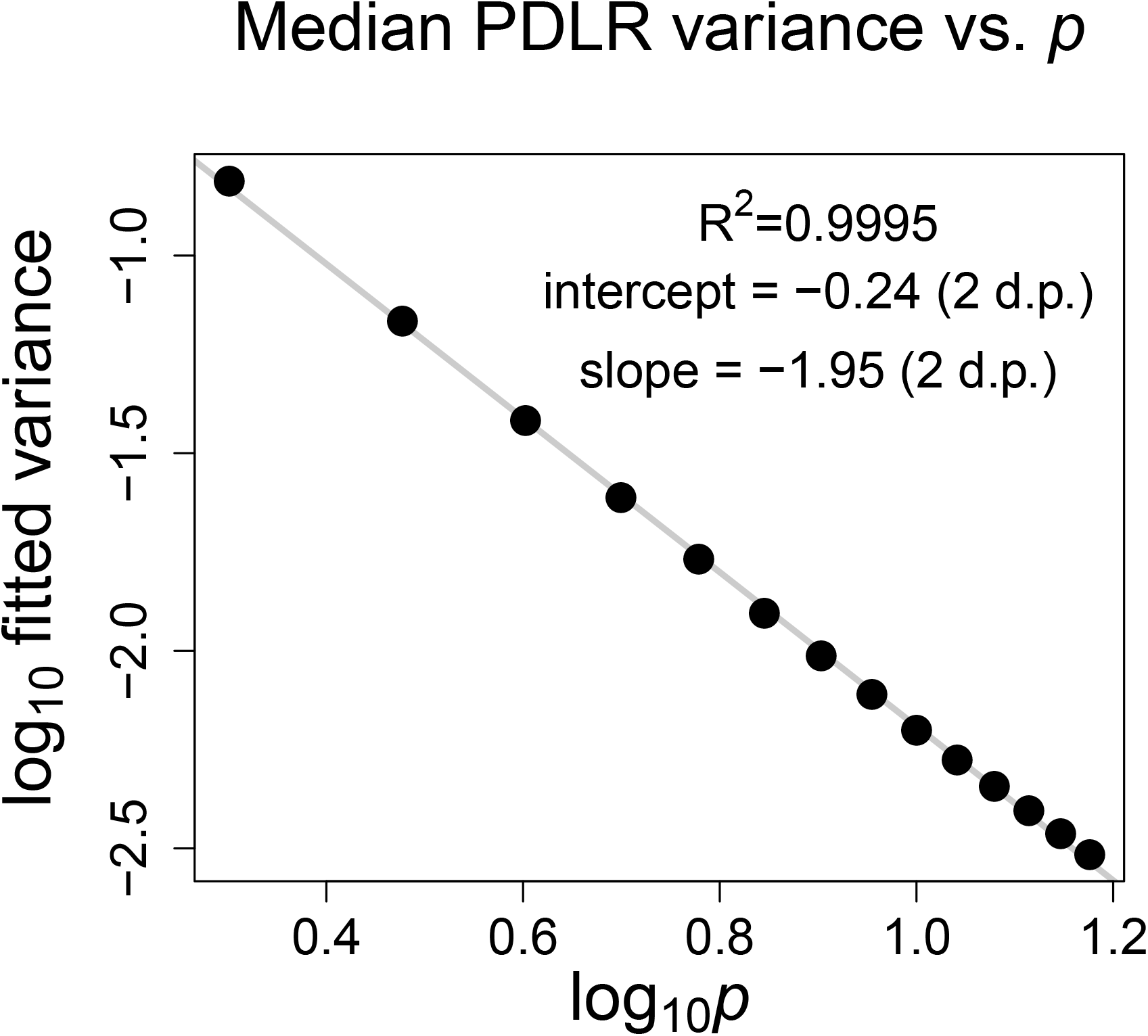
Following Figure 16, two sets of 10000 points are drawn from a *p*-dimensional normal distribution with *p* = 2, 3, …, 15 with mean of **0**, variance of one and correlation of zero. The PDLR distribution is calculated between the two sets of points and a Gaussian model is fit with one peak, and the variance of the distribution is extracted. This is repeated 100 times for each value of *p*, the median variance is obtained and plotted against *p*. The log-log plot shows a near perfect linear trend.

**Figure 18:**
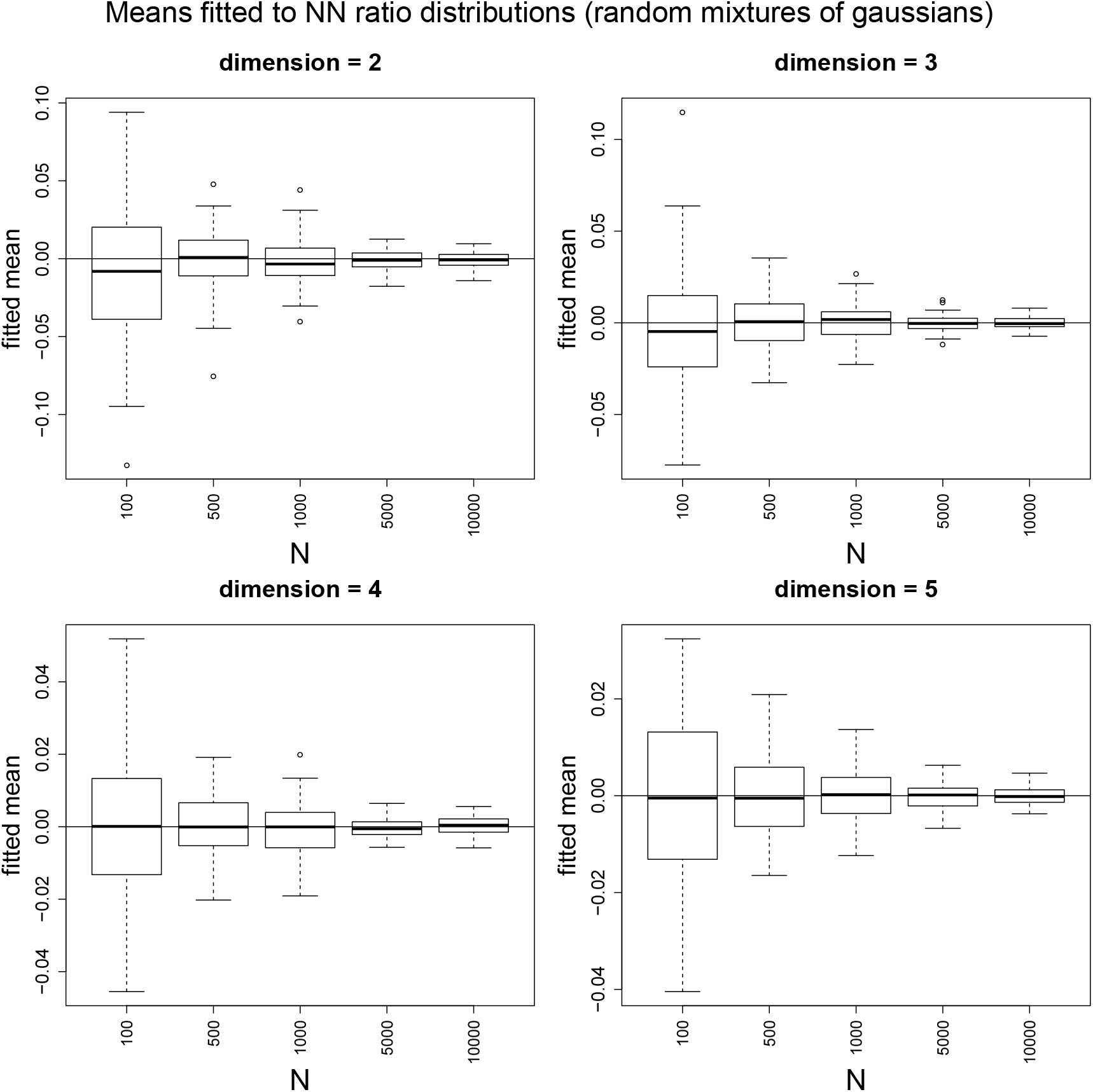
Two sets of *N* points are drawn from a *p*-dimensional random mixture of gaussians. The paired density ratio is calculated and fit using a single gaussian, and this is repeated 100 times for each combinations of *p* and *N*, with a unique mixture in each replicate. The fitted means are displayed as a boxplot over all the parameters. Similarly to Figure 15, the fitted means tend towards zero with increasingly smaller variance as *N* increases.

**Figure 19:**
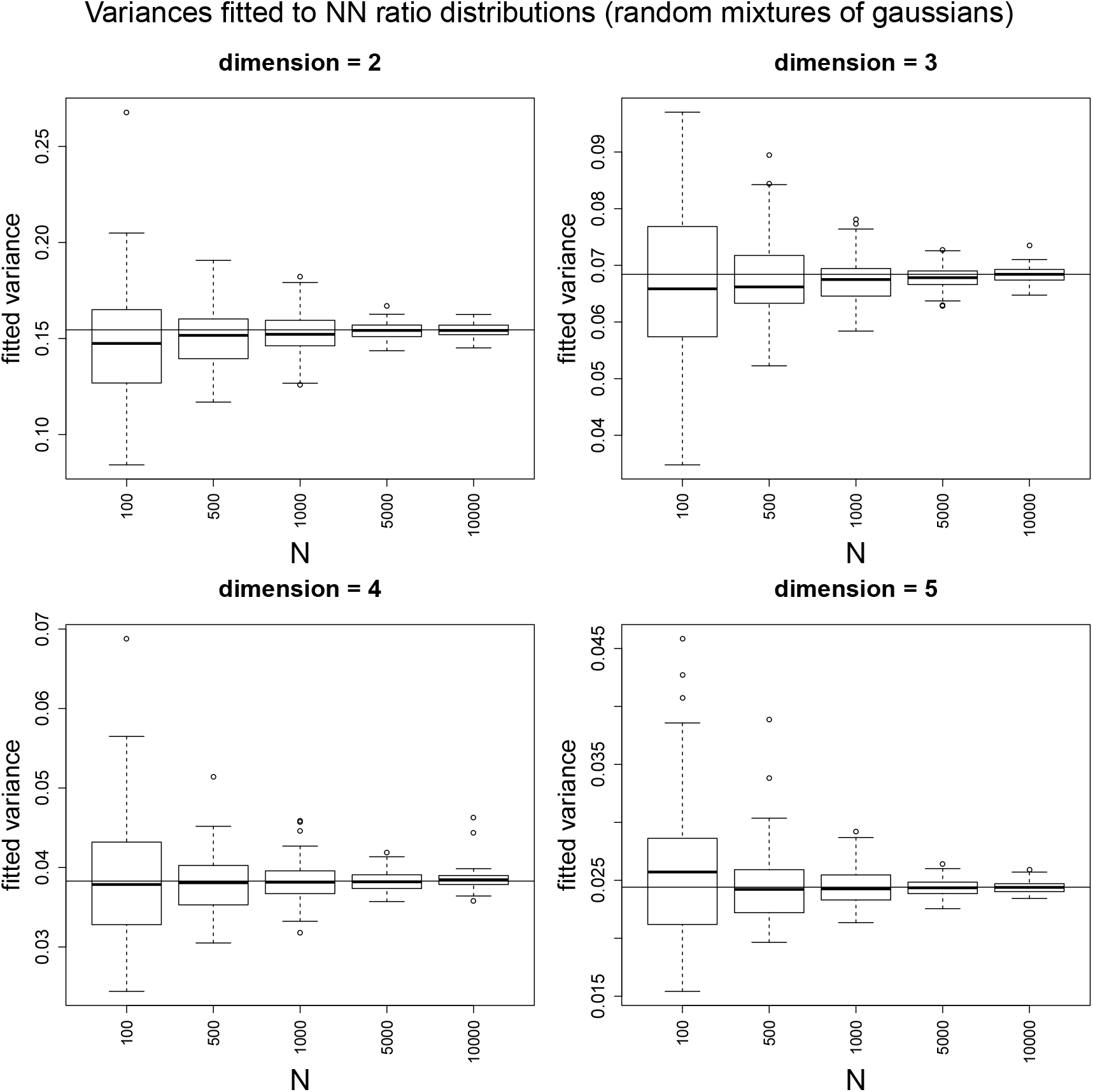
Two sets of *N* points are drawn from a *p*-dimensional random mixture of gaussians. The paired density ratio is calculated and fit using a single gaussian, and this is repeated 100 times for each combinations of *p* and *N*, with a unique mixture in each replicate. The fitted variances are displayed as a boxplot over all the parameters. Similarly to Figure 16, for each value of *p*, the fitted variances tend towards a fixed value with increasingly smaller variance as *N* increases. This value decreases as *p* increases. The horizontal lines are obtained from the values in Figure 17, meaning the median fitted PDLR variances are in good agreement with those arising from unimodal Gaussian distributions. It suggests that the self-PDLR distribution is a universal property of any arbitrary *p*-dimensional distribution.

To the best of my knowledge this is a novel result, but I am happy to stand corrected as I don’t have a comprehensive overview of the theoretical statistics literature. If it is indeed a novel result, a more theoretically minded person is welcome to flesh out the details, as it is outside my expertise.

### 8.4 Benchmarking

In order to benchmark the performance of the DensityMorph quasi-distance, a series of increasingly complex simulations are performed. Firstly, the DensityMorph distance is calculated between a reference *p*-dimensional normal point cloud (mean of **0**, correlation of zero, covariance of one) and one that differs with one of separation, covariance and correlation (**Figures 20, 21, 22**). Next the DensityMorph distance is calculated between a reference bimodal point cloud with points shared in a 1:9 ratio, and once that differs by ratio (**Figures 23**). The DensityMorph distances are compared to the Wasserstein distances. The DensityMorph distances roughly monotonically increase as the parameter changes, but not as smoothly as the Wasserstein distances, especially for small differences.

**Figure 20:**
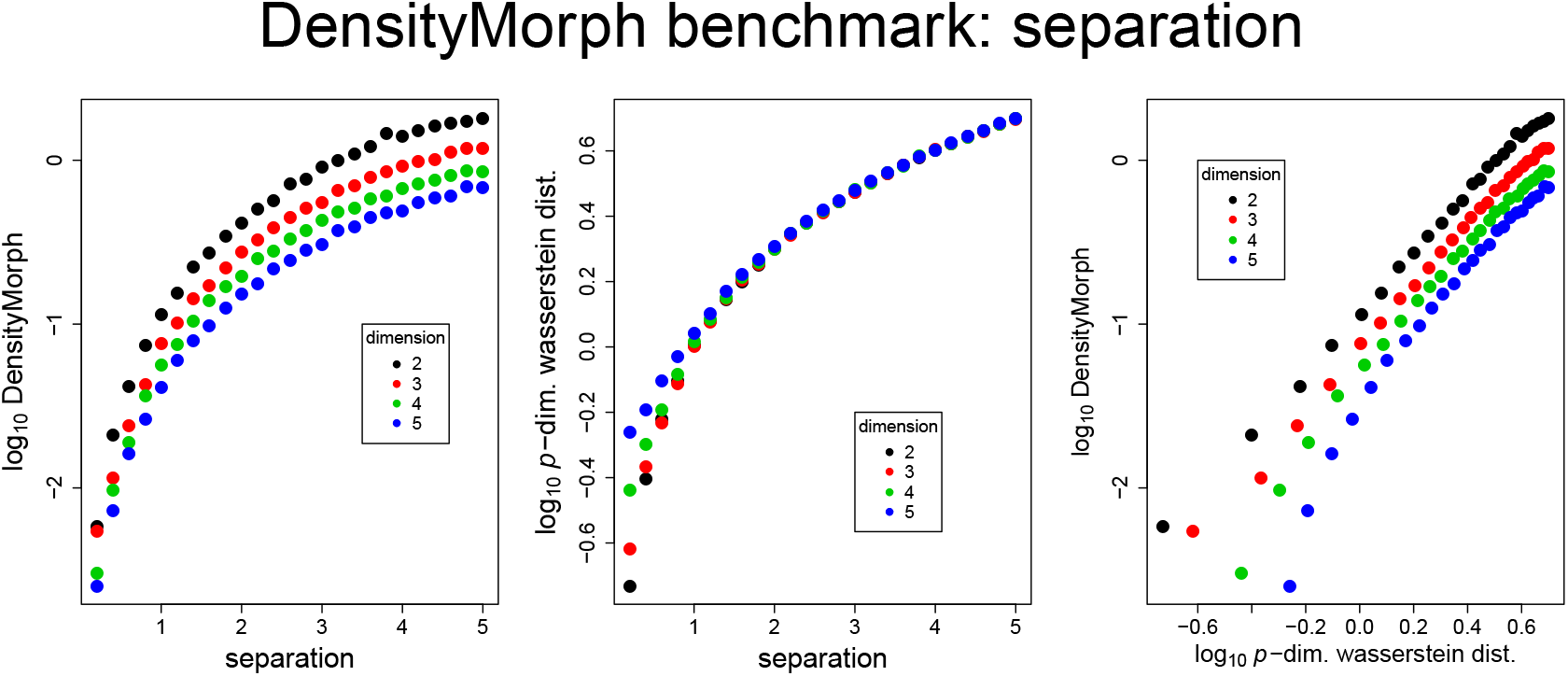
Benchmarking DensityMorph performance as a function of separation *d* between point clouds. For *p ∈ {*2, 3, 4, 5}, a point cloud *X*_1_ of mean **0**, variance of 1 and correlation of 0 is compared to point cloud *X*_2_(*c*) of mean (*d*, **0**), variance of 1 and correlation of 0 for *d ∈ {*0.2, 0.4, …, 5}. **Left** A plot of log_10_ *DM* (*X*_1_, *X*_2_(*c*)) vs. *c*, and *N*_1_ = *N*_2_ = 10000. **Middle** A plot of log_10_ *WD*(*X*_1_, *X*_2_(*c*)) vs. *c*, and *N*_1_ = *N*_2_ = 10000. **Right** A plot of log_10_ *DM* (*X*_1_, *X*_2_(*c*)) vs. log_10_ *WD*(*X*_1_, *X*_2_(*d*)).

**Figure 21:**
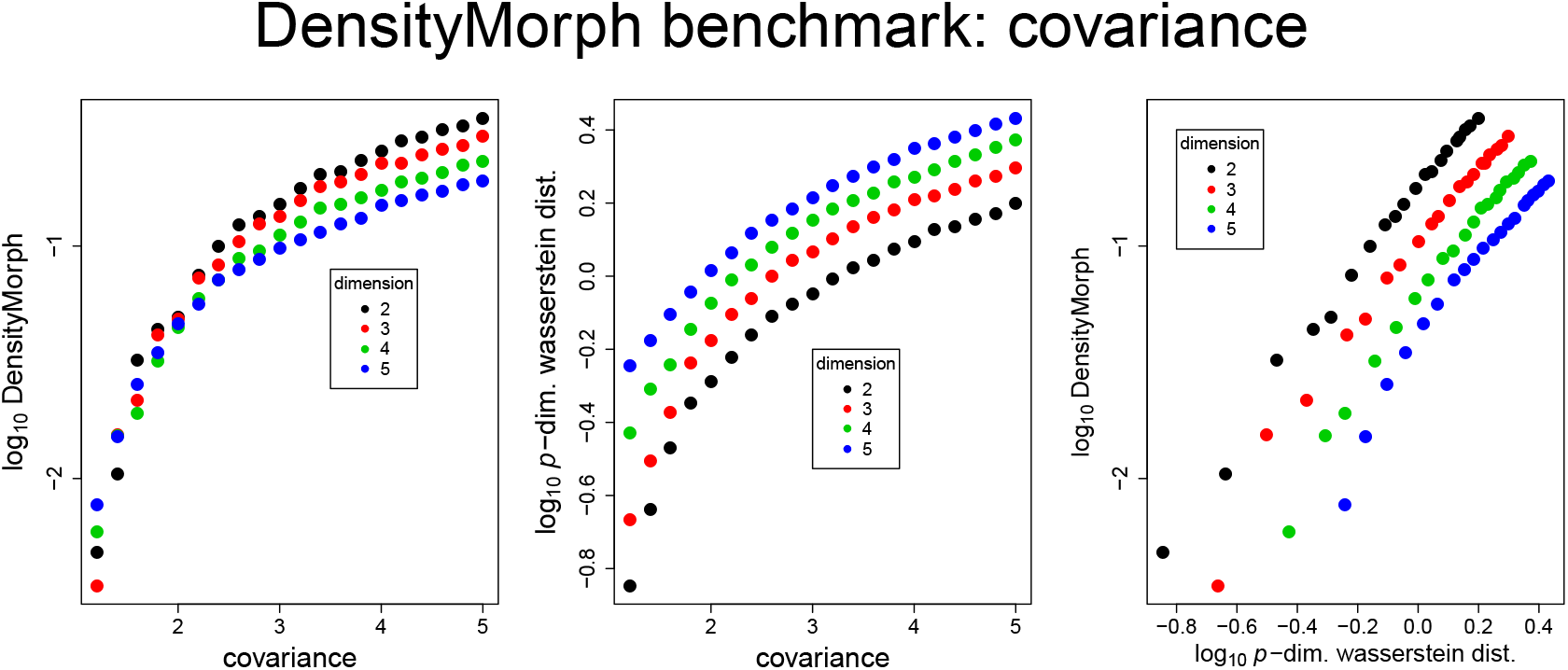
Benchmarking DensityMorph performance as a function of increasing covariance *c*. For *p ∈ {*2, 3, 4, 5}, a point cloud *X*_1_ of mean **0**, variance of 1 and correlation of 0 is compared to point cloud *X*_2_(*c*) of mean **0**, variance of *c* and correlation of 0 for *c ∈ {*1.2, 1.4, …, 5}. **Left** A plot of log_10_ *DM* (*X*_1_, *X*_2_(*c*)) vs. *c*, and *N*_1_ = *N*_2_ = 10000. **Middle** A plot of log_10_ *WD*(*X*_1_, *X*_2_(*c*)) vs. *c*, and *N*_1_ = *N*_2_ = 10000. **Right** A plot of log_10_ *DM* (*X*_1_, *X*_2_(*c*)) vs. log_10_ *WD*(*X*_1_, *X*_2_(*c*)).

**Figure 22:**
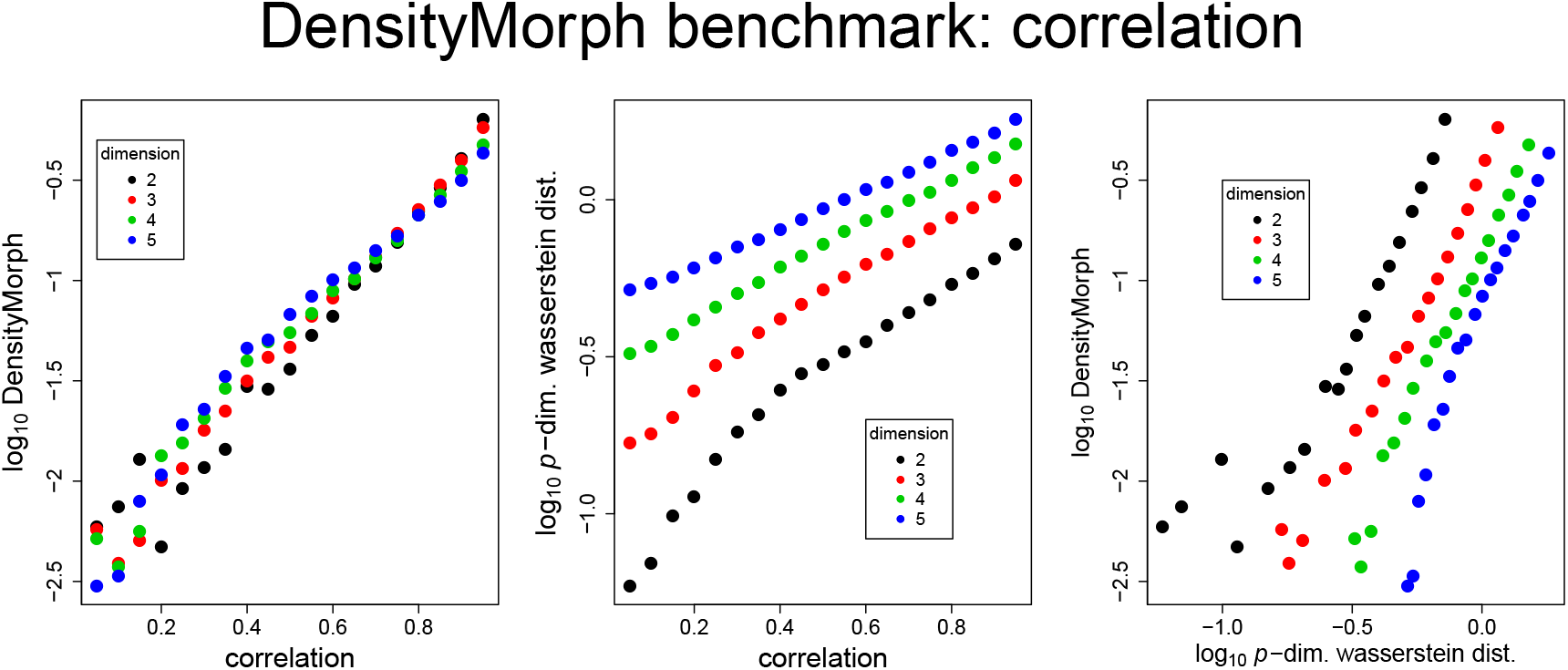
Benchmarking DensityMorph performance as a function of increasing correlation *r*. For *p ∈ {*2, 3, 4, 5}, a point cloud *X*_1_ of mean **0**, variance of 1 and correlation of 0 is compared to point cloud *X*_2_(*r*) of mean **0**, variance of 1 and correlation of *r* for *r ∈ {*0.05, 0.1, …, 0.95}. **Left** A plot of log_10_ *DM* (*X*_1_, *X*_2_(*r*)) vs. *r*, and *N*_1_ = *N*_2_ = 10000. **Middle** A plot of log_10_ *WD*(*X*_1_, *X*_2_(*r*)) vs. *r*, and *N*_1_ = *N*_2_ = 10000. **Right** A plot of log_10_ *DM* (*X*_1_, *X*_2_(*r*)) vs. log_10_ *WD*(*X*_1_, *X*_2_(*r*)).

**Figure 23:**
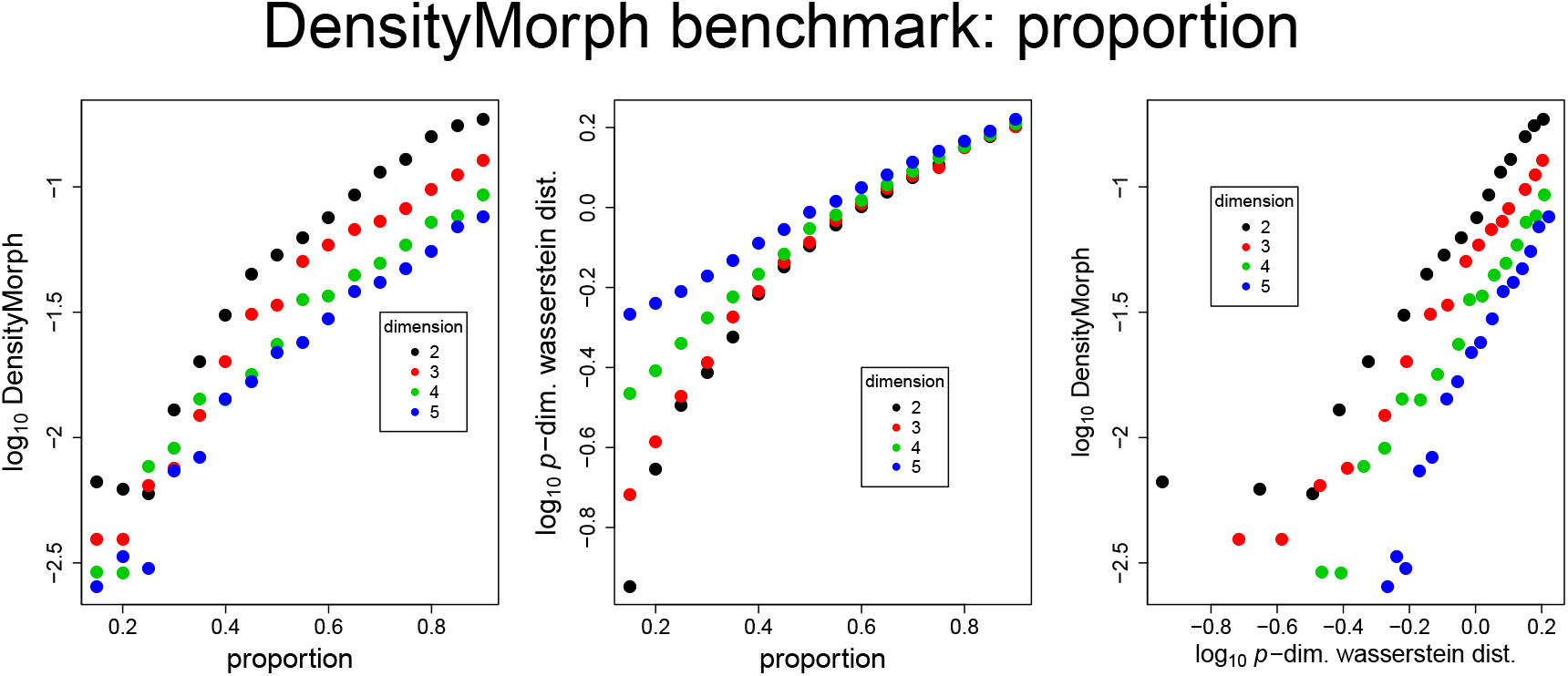
Benchmarking DensityMorph performance on bimodal point clouds as a function of proportion *b*. For *p ∈ {*2, 3, 4, 5}, a bimodal point cloud *X*_1_ of means **0** (10%) and (2, **0**) (90%), variance of 1, correlation of 0 is compared to a bimodal point cloud *X*_2_(*b*) of means **0** (*b*%) and (2, **0**) (100 *− b*%), variance of 1 and correlation of 0 for *b ∈ {*15, 20, …, 90}. **Left** A plot of log_10_ *DM* (*X*_1_, *X*_2_(*b*)) vs. *b*, and *N*_1_ = *N*_2_ = 10000. **Middle** A plot of log_10_ *WD*(*X*_1_, *X*_2_(*b*)) vs. *b*, and *N*_1_ = *N*_2_ = 10000. **Right** A plot of log_10_ *DM* (*X*_1_, *X*_2_(*b*)) vs. log_10_ *WD*(*X*_1_, *X*_2_(*b*)).

Next, *p*-dimensional normal point clouds are generated on a parameter grid with mean *∈ {*0, 1, …, 5}, covariance *∈ {*1, 2, …, 5} and correlation *∈ {*0.2, 0.4, 0.6, 0.8} (120 point clouds in total). DensityMorph is used to construct a 120 *×* 120 distance matrix, and principal *coordinates* analysis is used to embed the point clouds in a latent space. This is done three times, where the distance matrix is transformed element-wise to the power of *k ∈ {*0.1, 0.5, 1}. The latent spaces are plotted three times each, colour coded according to the parameters (**Figure 24**). The arrangement is globally driven by changes in mean and covariance, and locally by changes in correlation. For all three values of *k*, the arrangement is consistent with the parameter grid, and *k* = 0.5 is used in **Section 4**.

**Figure 24:**
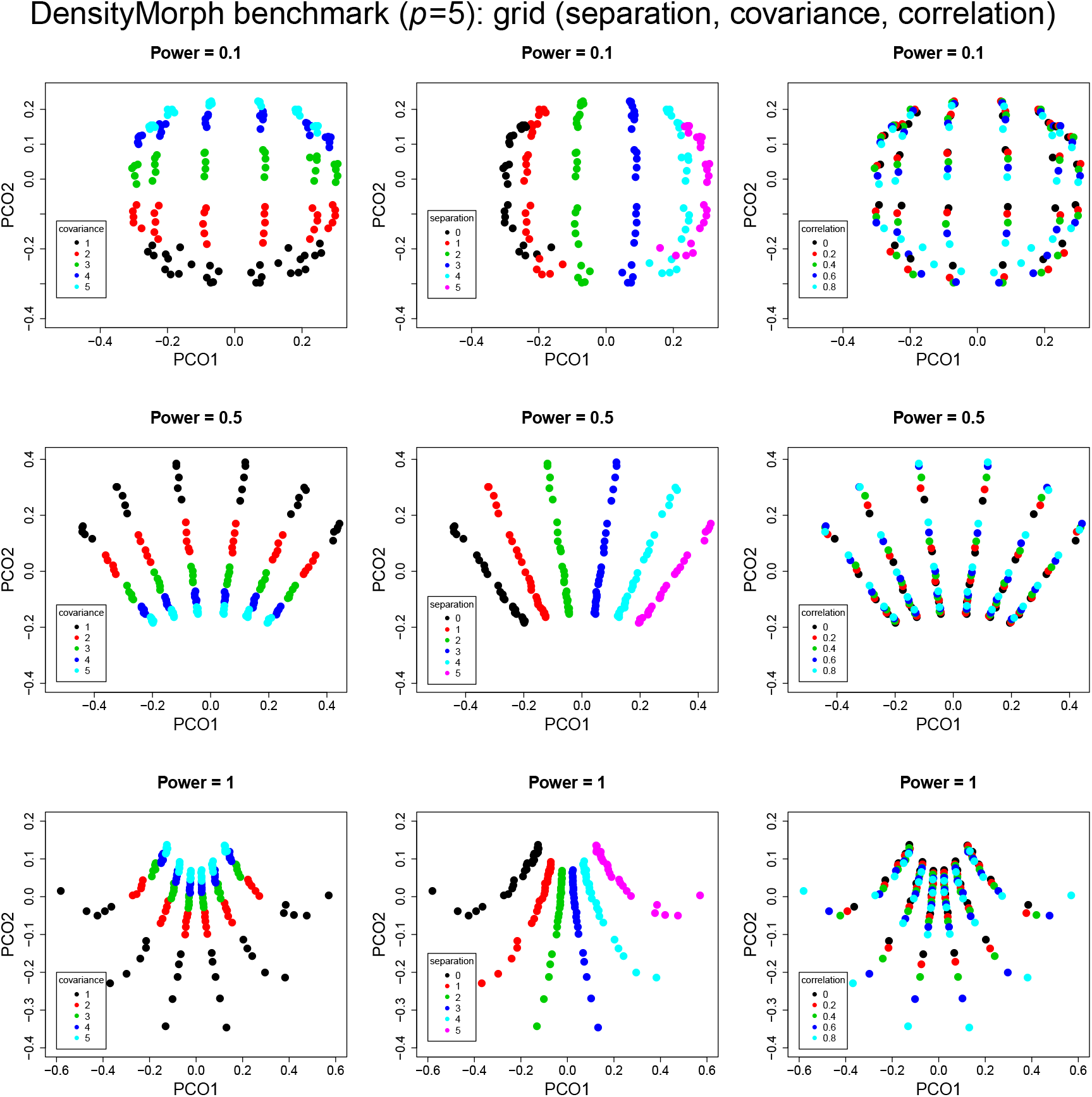
Benchmarking DensityMorph performance on a parameter grid for *p* = 5. For every combination of mean *M ∈ {*0, 1, …, 5}, covariance *c ∈ {*1, 2, …, 5} and correlation *r ∈ {*0.2, 0.4, 0.6, 0.8}, a point cloud is constructed and DensityMorph is used to construct a distance matrix. The distance matrix is transformed element-wise to the power of *k ∈ {*0.1, 0.5, 1} and principal *coordinate* analysis is performed to yield the latent space (not to be confused with principal component analysis). For each value of *k*, the point cloud is plotted three times, each colour coded according to *M, c* and *r*.

The final benchmark is to calculate a distance matrix with DensityMorph and Wasserstein distance over a set of point clouds arising from random mixtures of gaussians. Fifty point clouds of 10000 points are created and each underlying gaussian mixture is unique. As it is too computationally intensive to calculate the full Wasserstein distance matrix, 1000 points are sampled from each point cloud and the distances are calculated, and this is repeated 10 time and the mean distance is used. The DensityMorph distance matrix is calculated and there is a correlation of about 0.85 between two distance types (**Figure 25**). A procrustes analysis is performed with the two distance matrices and there is good agreement between the two ordinations (**Figure 26**).

**Figure 25:**
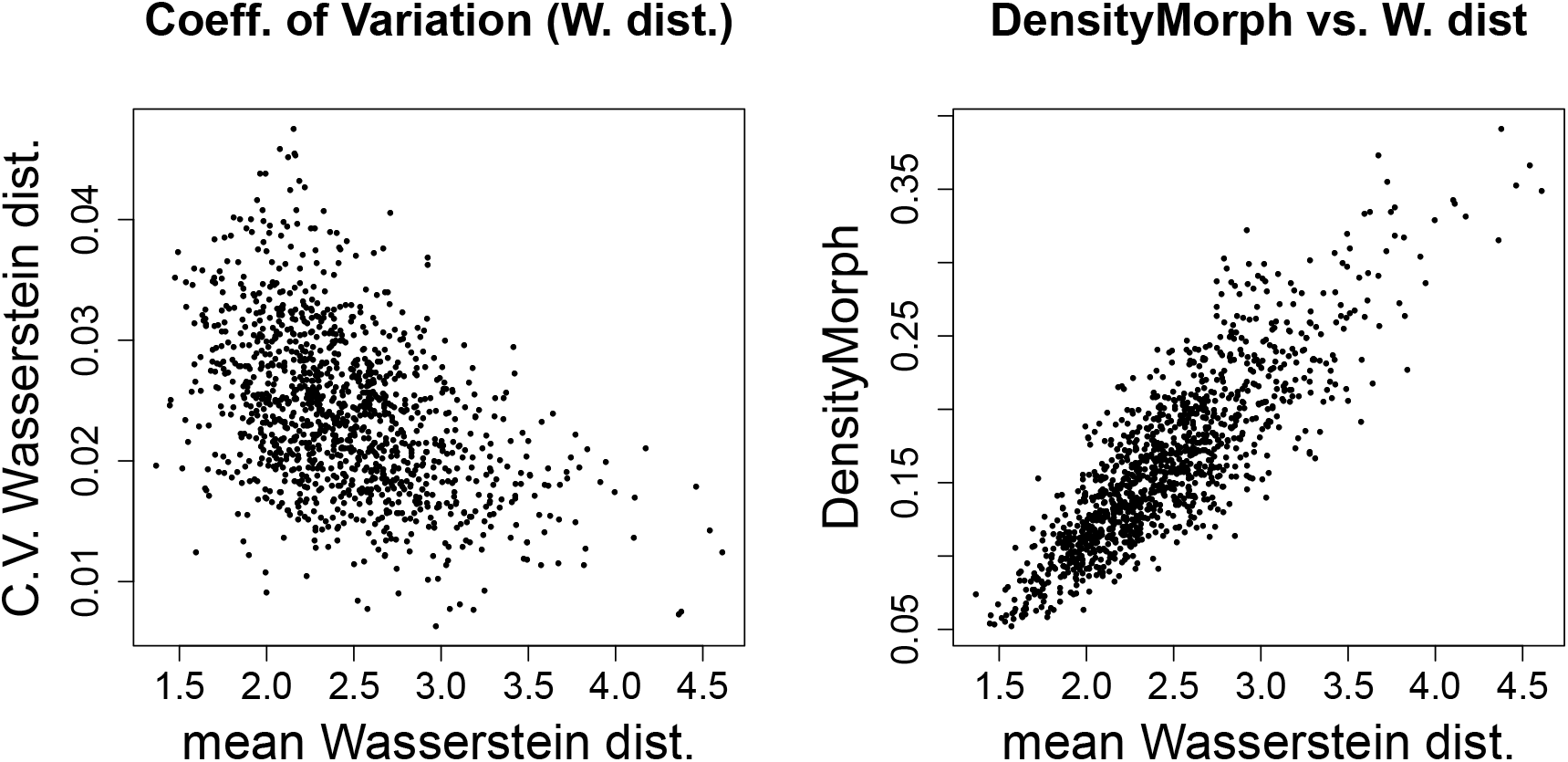
Benchmarking DensityMorph with a set of 50 arbitrary point clouds. Each point cloud is generated by superposing 5 subclouds of randomly selected mean on the interval (0, 5), randomly selected variance on the interval (1, 3), randomly selected correlation on the interval (*−*0.3, 0.3). The number of points in each subcloud is selected using a multinomial distribution with randomly selected probabilities, such that the total number of points is 10000. **Left** As it is too computationally intensive to calculate the full Wasserstein distance matrix, 1000 points are subsampled from each point cloud in order to calculate the Wasserstein distance matrix. This is repeated 10 times, and for each position in the distance matrix, the mean and coefficient of variation (CV) is calculated and plotted to show the variation. **Right** The DensityMorph distance matrix is calculated, and all distances are plotted against the corresponding mean Wasserstein distances. The correlation between the two quantities is 0.85 (2 d.p.).

**Figure 26:**
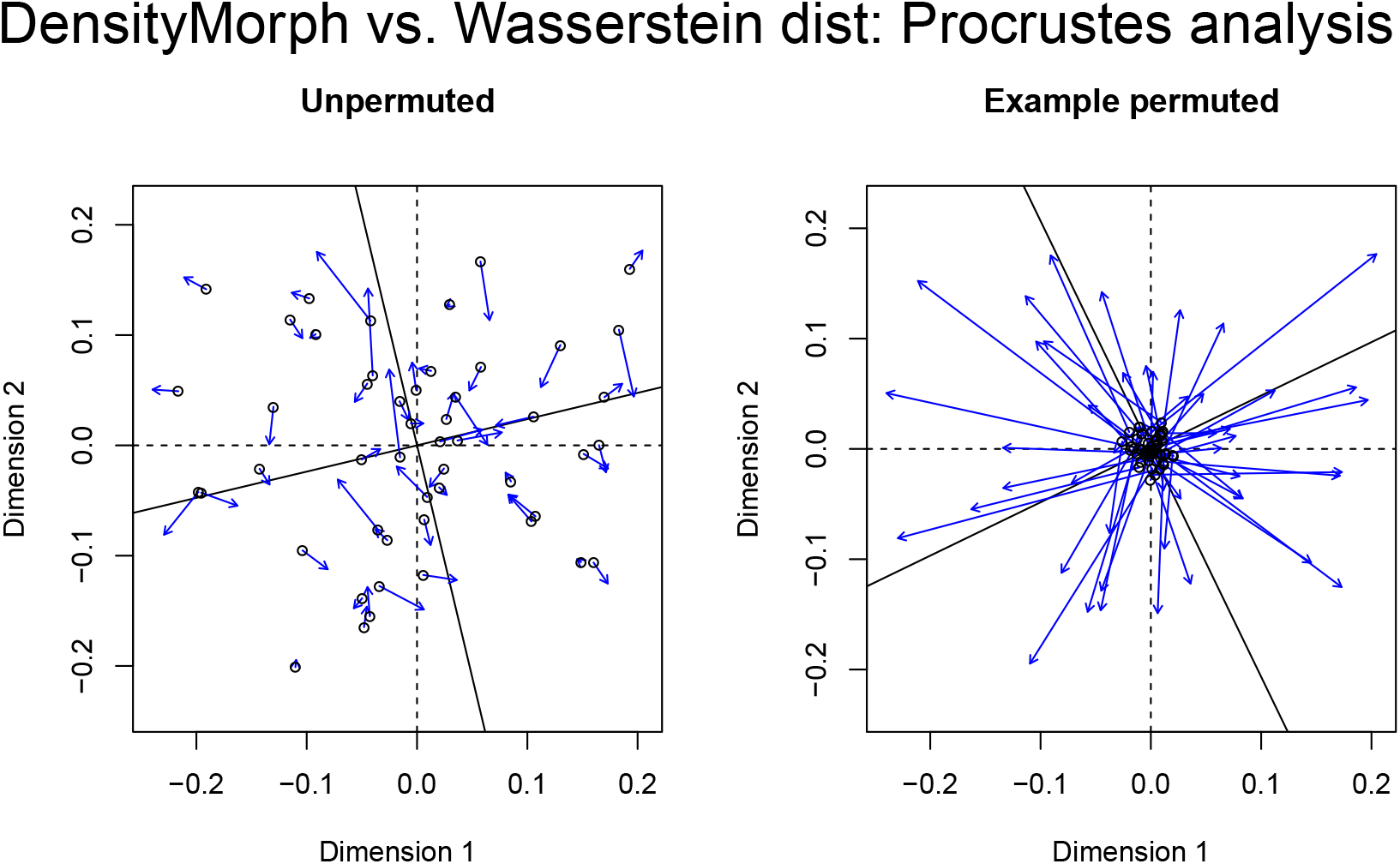
Benchmarking DensityMorph with a set of 50 arbitrary point clouds. **Left** The similarity between the DensityMorph distance matrix (transformed element-wise with *k* = 0.5) and the mean Wasserstein distance matrix is visualised using Procrustes analysis. **Right** The similarity between the permuted DensityMorph distance matrix (transformed element-wise with *k* = 0.5) and the permuted mean Wasserstein distance matrix is visualised using Procrustes analysis (row/column labels undergo the same permutation so the matrix structure remains the same).

## References

[1] T. Hastie, R. Tibshirani and J. Friedman (2009), Elements of Statistical Learning, Springer, New York, 2nd Edition

[2] V.M. Panaretos and Y. Zemel (2019), Statistical Aspects of Wasserstein Distances, Ann. Rev. Stat. Appl. 6:405–31

[3] J.C. Gower (1966), Some distance properties of latent root and vector methods used in multivariate analysis, Biometrika, 53:325–338

[4] S. Wold et. al. (1984), The collinearity problem in linear regression - the partial least squares approach to generalised inverses, SIAM J. Sci. Stat. Comput, 5:735–743

[5] C. Lucas et. al. (2020), Longitudinal analyses reveal immunological misfiring in severe COVID-19, Nature, https://doi.org/10.1038/s41586-020-2588-y

[6] P.J. Lewi (1989), Spectral Map Analysis: factorial analysis of contrasts, especially from log ratios, Chemometrics and Intelligent Laboratory Systems, 5:105–116

[7] B.A. Pereira et. al. (2019), CAF Subpopulations: a new reservoir of stromal targets in pancreatic cancer, Trends in Cancer, https://doi.org/10.1016/j.trecan.2019.09.010

[8] N. Bonneel et. al. (2011), Displacement interpolation using Lagrangian mass transport, ACM Transactions on Graphics (SIGGRAPH ASIA 2011), 30:6

